# Type I interferon responses to ischemic injury begin in the bone marrow of mice and humans and depend on Tet2, Nrf2, and Irf3

**DOI:** 10.1101/765404

**Authors:** David M. Calcagno, Richard P. Ng, Avinash Toomu, Claire Zhang, Kenneth Huang, Aaron D. Aguirre, Ralph Weissleder, Lori B. Daniels, Zhenxing Fu, Kevin R. King

## Abstract

Sterile tissue injury locally activates innate immune responses via interactions with damage associated molecular patterns (DAMPs). Here, by analyzing ∼120K single cell transcriptomes after myocardial infarction (MI) in mice and humans, we show neutrophil and monocyte subsets induce type I interferon (IFN) stimulated genes (ISGs) in myeloid progenitors of the bone marrow, far from the site of injury. In patients with acute MI, peripheral blood neutrophils and monocytes express ISGs at levels far beyond healthy individuals and comparable to patients with lupus. In the bone marrow of Tet2^-/-^ mice, ISGs are spontaneously induced in myeloid progenitors and their progeny. In the heart, IFN responses are negatively regulated by Ccr2- resident macrophages in a Nrf2-dependent fashion. Our results show post-MI IFN signaling begins in the bone marrow, implicate multiple transcription factors in its regulation (Tet2, Irf3, Nrf2), and provide a clinical biomarker (ISG score) for studying post-MI IFN signaling in patients.

## Introduction

Ischemic tissue injury is the initiating event underlying the most common causes of death in the world(Collaborators, 2018). Acute ischemia in the heart, known as myocardial infarction (MI), provokes an emergency myelopoietic response in the bone marrow that rapidly increases production of neutrophils and monocytes, and leads to peripheral blood leukocytosis, tissue infiltration, and organ dysfunction (e.g. heart failure), the hallmarks of acute inflammation (Boettcher and Manz, 2016; Dutta et al., 2015; Herault et al., 2017; Hoyer et al., 2019; Manz and Boettcher, 2014; Nahrendorf et al., 2007). While there is consensus that emergency myelopoiesis increases the *number* of circulating myeloid cells, it is unknown whether emergency cells are *functionally* distinct from their non-emergency counterparts.

At the site of injury, myeloid cells infiltrate tissue as overlapping waves of neutrophils and monocytes. Neutrophils, which peak at post-MI days 1-2, generate reactive oxygen species, elaborate protease- and myeloperoxidase-containing granules, and are thought to exacerbate tissue damage (Vinten-Johansen, 2004). Although protective neutrophil subsets have also been proposed, the full functional diversity of infarct neutrophils remains largely unexplored (Horckmans et al., 2017; Puhl and Steffens, 2019). Monocytes, which peak at post-MI days 3-4, infiltrate and differentiate into functionally heterogeneous Ccr2+ macrophage subsets with both proinflammatory and reparative phenotypes (Granger and Korthuis, 1995; Leuschner et al., 2012; Nahrendorf et al., 2007; Vinten-Johansen, 2004). Also present within the infarct are Ccr2- resident macrophages, which are proposed to play protective roles by incompletely understood mechanisms {Epelman et al. 2014; Dick et al. 2019; Bajpai, 2019 #8815}. Broadly speaking, myeloid cells are thought to develop specialized effector functions as a consequence of interactions with damage associated molecular patterns (DAMPs), cytokines, and other stimuli within the injured tissue microenvironment (Arslan et al., 2011; Kaczmarek et al., 2013). Here, we examine whether innate immune pathways may be activated remotely, prior to infiltrating the infarcted heart.

The type I IFN response is an innate immune pathway best known for its roles in the anti-viral response and its association with auto-inflammatory diseases such as lupus (Furie et al., 2017; Lee-Kirsch, 2017; Muller et al., 1994). We and others recently discovered that pathologic type I IFN responses also occur after ischemic injury in the heart. We showed that genetic or pharmacologic inhibition of IFN signaling reduced inflammation, limited adverse ventricular remodeling, and improved survival (Cao et al., 2018; King et al., 2017). Circumstantial evidence localized the response to monocyte-derived macrophages within the infarcted heart, and as a consequence, it was thought to be unconfirmable in patients because it is unsafe to biopsy the recently infarcted human heart. However, our data did not completely rule out activation in the blood or bone marrow prior to arrival at the heart. Here, we hypothesize that peripheral blood myeloid cells may exhibit functional heterogeneity prior to entering the heart, and that it went obscured in previous studies to the use of ensemble measurement techniques (e.g. qPCR and bulk RNA Seq). To test this hypothesis, we performed single cell RNA-Seq analysis of the bone marrow, blood, and heart after MI (Zilionis et al., 2017) and defined the origins, regulation, and human relevance of MI-induced type I IFN signaling.

Our results show that MI-induced IFN responses begin remotely, in the distant bone marrow, at the origins of emergency myelopoiesis. We analyzed >120,000 single cell transcriptomes from the hearts, blood, and bone marrow of mice and humans and combined them with integrated re-analysis of an additional 64,043 published transcriptomes to define the intracardiac states, trajectories, and inducible programs of post-MI neutrophils and monocytes within multiple tissue compartments across time. In patients presenting with acute MI, we observed defined subsets of human peripheral blood neutrophils and monocytes expressing ISGs at levels far beyond healthy subjects and comparable to patients with lupus. In mice, we confirm these findings and show that ISG expression is regulated by Tet2 in the bone marrow and Nrf2 in resident macrophages of the heart. This data provides the first comprehensive single cell transcriptomic maps of murine neutrophils, monocytes, and macrophages of the heart, blood, and bone marrow on days 1-4 after MI.

## Results

### IFN induced neutrophils and monocytes are elevated in peripheral blood of humans after MI

Healthy human peripheral blood monocytes have been extensively characterized using single cell transcriptomics; however, the changes that accompany sterile tissue injuries such as MI remain unknown. To understand human myeloid responses to sterile injury with single cell resolution, we collected peripheral blood from a patient presenting 48 hours after onset of chest pain with rising serum cardiac troponin levels signifying a non-ST elevation MI (NSTEMI). Rather than using commercial research collection tubes that isolate peripheral blood monocytes (PBMCs) but discard neutrophils, we collected whole blood, performed red blood cell lysis, and resuspended all of the remaining cells for flow sorting. Cells were stained with DAPI and cell surface CD235a antibody, flow sorted to purify live non-doublet non-RBCs, and barcoded using commercial 10X Genomics single cell RNA-Seq workflows. We integrated the results with 64,043 previously reported single cell transcriptomes from peripheral blood monocytes (PBMCs) of healthy individuals (Figure 1A). We then subset CD14+ monocytes and performed sub-clustering which revealed five clusters, four of which had comparable representation from healthy and post-MI cells (Figure 1B) and one that was dominated by cells from the post-MI sample (96.4%) (Figure 1C) with only rare representation by cells from the healthy samples (3.6%) (Figure 1D). Differential gene expression comparing the MI-enriched cluster to all other monocytes revealed a marker gene list almost exclusively comprised of previously annotated ISGs, suggesting that post-MI monocytes are IFN-induced before they arrive at the infarcted heart (Figure 1E). We summed expression levels of the top ISGs to create a composite ISG score (see methods) which was markedly elevated in the post-MI monocyte cluster (Figure 1F). Quantitative comparisons of ISG scores showed that post-MI monocytes were significantly elevated (9.56 ± .15) compared to healthy PBMCs (7.08 ± .06), (*P*<0.0001). For comparison, we independently analyzed published single cell transcriptomes from PBMCs of 8 patients with the IFN-associated autoimmune disease systemic lupus erythematous (SLE) and measured average ISG scores only modestly above those of post-MI monocytes (10.26 ± .15) (*P*<0.05) (Figure 1G).

**Figure 1.**
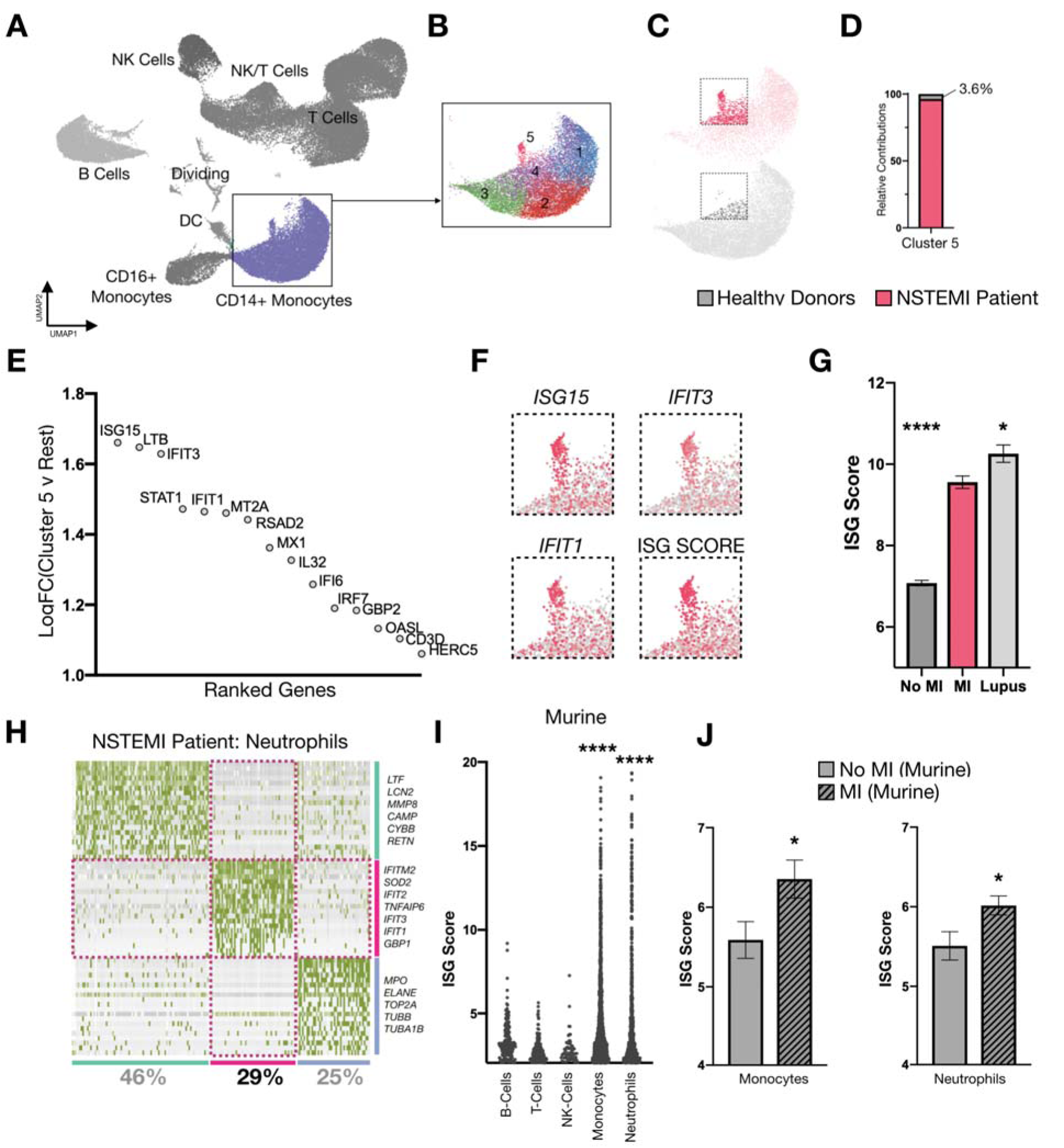
IFN induced neutrophils and monocytes are elevated in peripheral blood of humans and mice after acute MI. (**A-H**) Single cell RNA-seq data of peripheral blood mononuclear cells from healthy (n= 7 patients, n = 64,043 cells) and NSTEMI patients (n = 6,487 cells; 714 ng/L Troponin T Gen 5 at 12 hrs post onset-time; blood collected 28 hrs post onset-time). (**A-C**) UMAP of (**A**) all peripheral blood mononuclear cells and (**B**) clustered CD14+ monocytes in (**C**) healthy and NSTEMI patients. (**D**) Relative contributions of healthy and NSTEMI patients to cluster 5 monocytes, normalized by total quantity of CD14+ monocytes in each sample. (**E**) Top 15 upregulated genes in cluster 5 compared to clusters 1-4 using Wilcoxon Rank Sums test. All genes shown had significant (< .001) Bonferroni corrected p-values. (**F**) Feature plot highlighting select IFN stimulated genes (ISGs) and ISG score (defined as summed expression of top ISGs, see methods) in cluster 5. (**G**) ISG score in healthy, NSTEMI and Lupus patients. (**H**) Heatmap of neutrophils bioinformatically isolated from NSTEMI patient and subclustered to reveal ISG+ subset. (**I**) ISG scores of murine peripheral blood cells after MI, D1-D3 combined (6 mice; 25,469 cells). (**J**) ISG scores of monocytes and neutrophils at steady state and after MI (D1-D3). Data are shown as mean ± s.e.m. * P < .05, **** P< .0001, Mann-Whitney test.

We next bioinformatically isolated neutrophils from the post-MI peripheral blood samples. Little is known about single cell transcriptomes of human peripheral blood neutrophils because they are commonly discarded by commercial clinical collection tubes. Unbiased sub-clustering of post-MI neutrophils revealed IFN-induced (ISG+) and uninduced (ISG-) populations that spanned the spectrum of maturity (Figure 1H). Taken together, these results suggest that human peripheral blood contains ISG+ IFN induced cells (IFNICs) and ISG- uninduced cells of both neutrophil and monocyte myeloid lineages after MI.

### IFN induced neutrophils and monocytes are elevated in peripheral blood of mice after MI

To explore the origins, differentiation patterns, and regulation of myeloid IFN responses, we ligated the coronary arteries of mice to induce MI and performed single cell RNA-Seq on post-MI days 1-4 focusing on three compartments – the bone marrow, where myeloid cells originate and differentiate from progenitors; the blood, through which they traffic; and the infarcted heart, where they infiltrate and differentiate into subsets with specialized functions. We first analyzed peripheral blood myeloid cells to determine if IFN signaling was induced by MI in mice. Consistent with our results in humans, single cell transcriptomes of neutrophils and monocytes were the dominant sources of ISG transcripts in mice, exhibiting significantly increased ISG scores compared to B cells, T cells, or NK cells (*P* < 0.0001) (Figure 1I). As in humans, we observed both ISG+ and ISG- neutrophils and monocytes in the post-MI blood of mice (Figure S1A-D). To test whether ISGs were increased after MI, we separately compared post-MI to no-MI ISG scores of monocytes and neutrophils and observed significant increases (Figure 1J). Taken together, these data suggest that MI induces a type I IFN response that is measurable by quantifying ISG+ and ISG- peripheral blood neutrophils and monocytes with single cell resolution in mice and humans.

### Neutrophils differentiate into mirrored ISG+ and ISG- neutrophil subsets within the infarcted mouse heart

We next directed our attention to the myeloid cells within the infarcted heart. We flushed and enzymatically digested infarcted mouse hearts, surface antibody stained and inclusively FACS sorted the released cells by gating only on live single non-erythrocytes (Dapi^LOW^, Ter119^LOW^) followed by microfluidic droplet-based single cell RNA-Seq using custom inDrop or 10X Genomics barcoding platforms (Table 1). Coarse clustering and bioinformatic subsetting of the resulting transcriptomes revealed a neutrophil-predominant distribution on day 1 and a monocyte-macrophage-predominant distribution on day 4 within the infarcted heart consistent with previously reported immunophenotyping and flow cytometry data (Figure S2A-D) (Nahrendorf et al., 2007). We bioinformatically isolated neutrophils and monocytes and separately reclustered them to investigate their post-MI intracardiac subsets.

**Table 1.** Summary of murine single-cell RNA seq experiments.

| Tissue | Days post-MI | Single Cell Platform | n = mice | Mouse | n = cells |
| --- | --- | --- | --- | --- | --- |
| Heart | 1,2,4 | inDrop | 3 | WT | 36,128 |
| Heart | 4 | inDrop | 1 | IFNAR<sup>-/-</sup> | 1,417 |
| Heart | 4 | inDrop | 1 | Irf3<sup>-/-</sup> | 3,227 |
| Heart | 4 | 10X | 2 | IFNAR Ab | 8,329 |
| Heart | 4 | 10X | 2 | Nrf2<sup>-/-</sup> | 10,912 |
| Blood | 0,1,2,3 | 10X | 2 | WT | 25,469 |
| Bone Marrow | 1,2,3 | 10X | 1 | WT | 6,539 |
| Bone Marrow | 0,1 | 10X | 1 | Tet2<sup>-/-</sup> | 23,868 |

Neutrophils are short-lived first responders to sterile tissue injuries such as MI. In contrast to monocytes and macrophages, little is known about how neutrophils diversify and specialize after entering the injured heart (Ng et al., 2019; Silvestre-Roig et al., 2016). We bioinformatically subset neutrophils (*S100a8, S100a9, Retnlg, Mmp9*, *Cxcr2*)^HI^, renormalized, scaled, and reclustered them to explore neutrophil heterogeneity. This revealed five intracardiac neutrophil subsets with distinct temporal patterns between D1 and D4 post-MI (Fig. 2A-C, Figure S3A-C, Table 2). The parent neutrophil subset (Hrt-N1), characterized by expression of *Retnlg*, was present in every sample at every time point and mirrored peripheral blood neutrophils (Figure 2A-C, Figure S3A-C, Table 3A). Hrt-N2 neutrophils expressed ISGs as early as day 1 after MI and as late as day 4 (Figure 2A-C, Figure S2B and S3A-C, Table 3B). This was unexpected since prior work had only uncovered ISG expression within monocyte-derived infarct macrophages (Cao et al., 2018; King et al., 2017). We termed these cells neutrophil IFN induced cells (nIFNICs). The Hrt-N3 subset expressed genes associated with NFκB activation, including *Nfkb1, Icam1, Il1a, Sod2, and Tnip1* (Figure 2A-C, Figure S3A-C, Table 3C) while the Hrt-N4 population expressed genes associated with *Hif-1α* activation, including *Egln3, Hilpda, and Vegfa* (Figure 2A-C, Figure S3A-C, Table 3D) (*Walmsley et al., 2011*). Both Hrt-N3 and Hrt-N4 were prominent on day 1 but declined by day 4 in concert with emergence of a new neutrophil subset (Hrt-N5), characterized by expression of *Siglecf* (Figure 2A-C, Figure S3A-C, Table 3E). Although *Siglecf* is best known as an eosinophil marker and is often gated against when flow sorting neutrophils, recent single cell transcriptomic profiling in the context of murine lung cancer identified a similar SiglecF-expressing neutrophil (Engblom et al., 2017; Zilionis et al., 2019). Indeed, we were able to confirm, using surface antibody staining, that neutrophils (Cd11b+, Ly6G+) are separable into Siglecf^HI^ and Siglecf^LOW^ subpopulations, which allowed FACS sorting and qPCR validation of the single cell RNA-Seq findings (Figure S4). These results demonstrate that neutrophils are functionally heterogeneous across time and that SiglecF neutrophils may play unrecognized roles within the infarct during late phases of the immune response when remodeling is thought to be dominated by macrophages.

**Figure 2.**
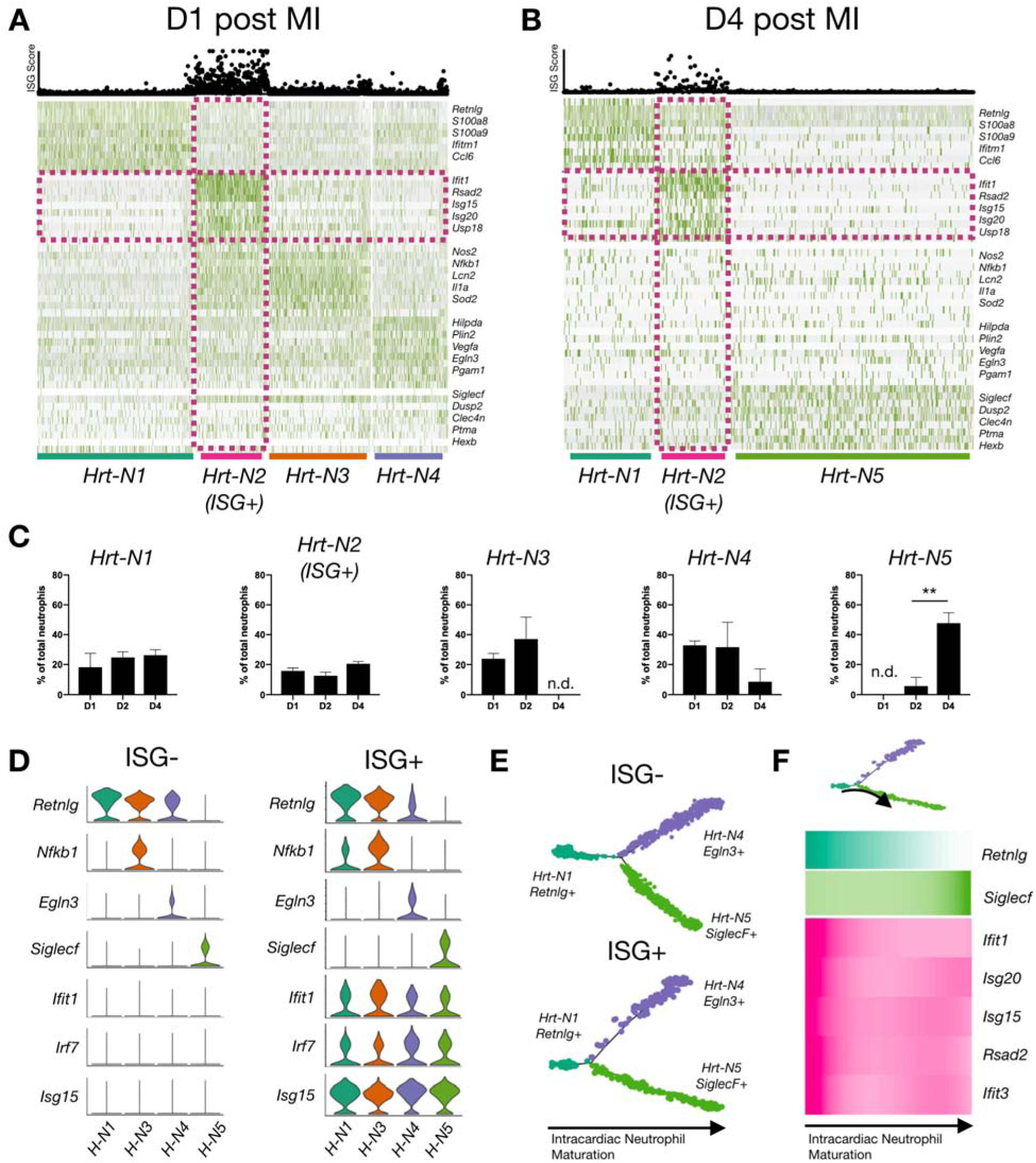
Neutrophils differentiate into mirrored ISG+ and ISG- neutrophil subsets within the infarcted mouse heart. (**A,B**) Representative heatmaps and ISG scores of intracardiac neutrophils on (**A**) days 1 (n = 2,285 cells, 1 mouse) and (**B**) day 4 (n = 720 cells, 1 mouse) after MI (ISG-expressing cells and marker genes outlined in pink). (**C**) Time course of intracardiac neutrophils subsets (n = 3 mice per day). Comparisons are non-significant unless noted. (**D**) Violin plots of subclustered ISG- and ISG+ cells (D1, D2 and D4 post-MI combined). Top marker gene for each neutrophil subset shown. (**E,F**) Pseudotime trajectory (Monocle) on ISG- and ISG+ (**E**) and heatmap of neutrophil maturation and ISG expression vs pseudotime (**F**) from Retnlg^HI^Siglecf^LOW^ to Retnlg^LOW^Siglecf^HI^. Data are shown as mean ± s.e.m. ** *P< 0.01,* unpaired two-sided student t-test.

**Table 2.** Quantification of intracardiac neutrophil subsets as a function of time post-MI by single cell RNA seq.

| Day post-MI | # of Hrt-N1 | # of Hrt-N2 | # of Hrt-N3 | # of Hrt-N4 | # of Hrt-N5 | Total Cells |
| --- | --- | --- | --- | --- | --- | --- |
| 1 | 620 | 441 | 745 | 682 | 0 | 2488 |
| 1 | 565 | 266 | 679 | 388 | 0 | 1898 |
| 1 | 208 | 133 | 0 | 173 | 0 | 514 |
| 2 | 195 | 222 | 392 | 232 | 0 | 1041 |
| 2 | 376 | 100 | 322 | 0 | 445 | 1243 |
| 2 | 254 | 156 | 547 | 0 | 0 | 957 |
| 4 | 214 | 163 | 0 | 0 | 343 | 720 |
| 4 | 127 | 100 | 0 | 0 | 339 | 566 |
| 4 | 123 | 96 | 0 | 368 | 123 | 710 |

**Table 3A.** Top 10 differentially expressed genes in intracardiac neutrophil subset Hrt-N1 relative to other neutrophils. Wilcoxon Rank Sums test.

| Gene | P-value | Avg. LogFC | % in Hrt-N1 | % in Rest | Adj. P-value |
| --- | --- | --- | --- | --- | --- |
| Anxa1 | 3.44E-128 | 1.6398562 | 0.634 | 0.171 | 4.33E-124 |
| Retnlg | 5.00E-123 | 1.0499896 | 0.982 | 0.847 | 6.30E-119 |
| Il1rn | 2.13E-121 | -1.7345475 | 0.398 | 0.823 | 2.68E-117 |
| S100a8 | 6.21E-117 | 0.8547638 | 0.994 | 0.937 | 7.82E-113 |
| Fth1 | 1.89E-114 | -0.5100967 | 0.998 | 1 | 2.38E-110 |
| Ccl3 | 7.99E-114 | -1.4544589 | 0.547 | 0.873 | 1.01E-109 |
| S100a9 | 1.45E-113 | 0.7566213 | 0.997 | 0.96 | 1.82E-109 |
| Cxcl2 | 1.15E-108 | -0.6155571 | 0.977 | 1 | 1.45E-104 |
| Ifitm6 | 2.36E-107 | 1.419944 | 0.629 | 0.197 | 2.97E-103 |
| Alox5ap | 3.45E-98 | 0.8571842 | 0.911 | 0.642 | 4.35E-94 |

**Table 3B.**
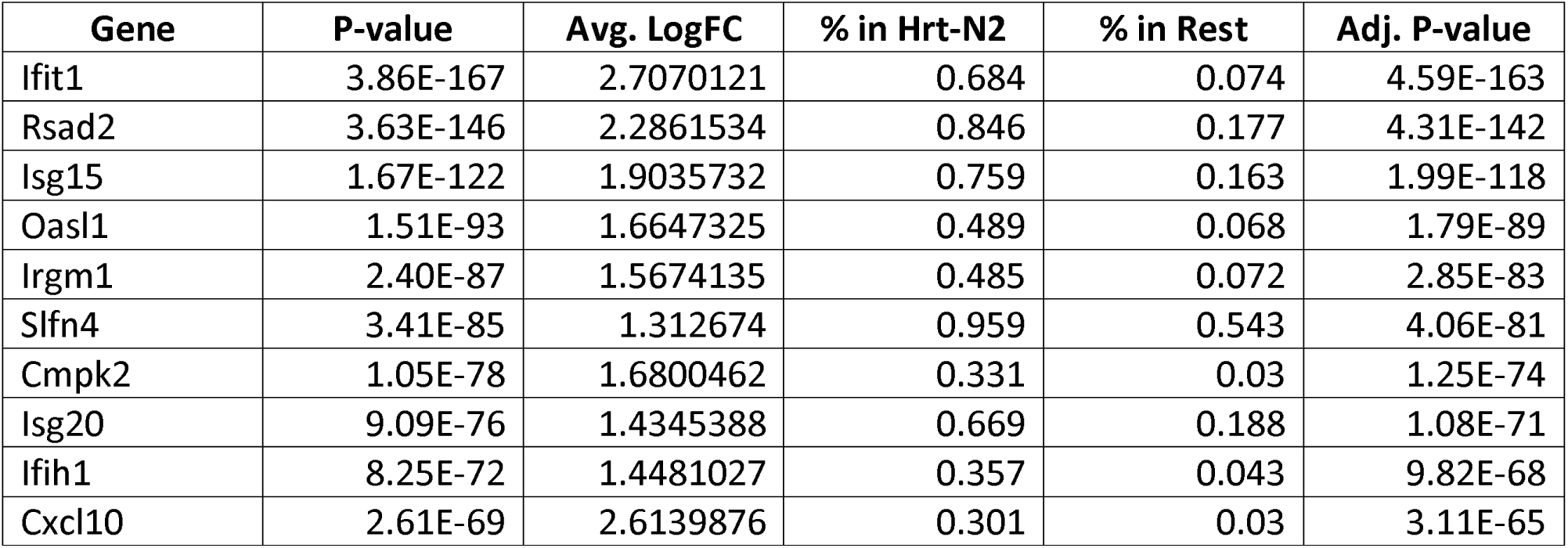
Top 10 differentially expressed genes in intracardiac neutrophil subset Hrt-N2. Wilcoxon Rank Sums test.

**Table 3C.**
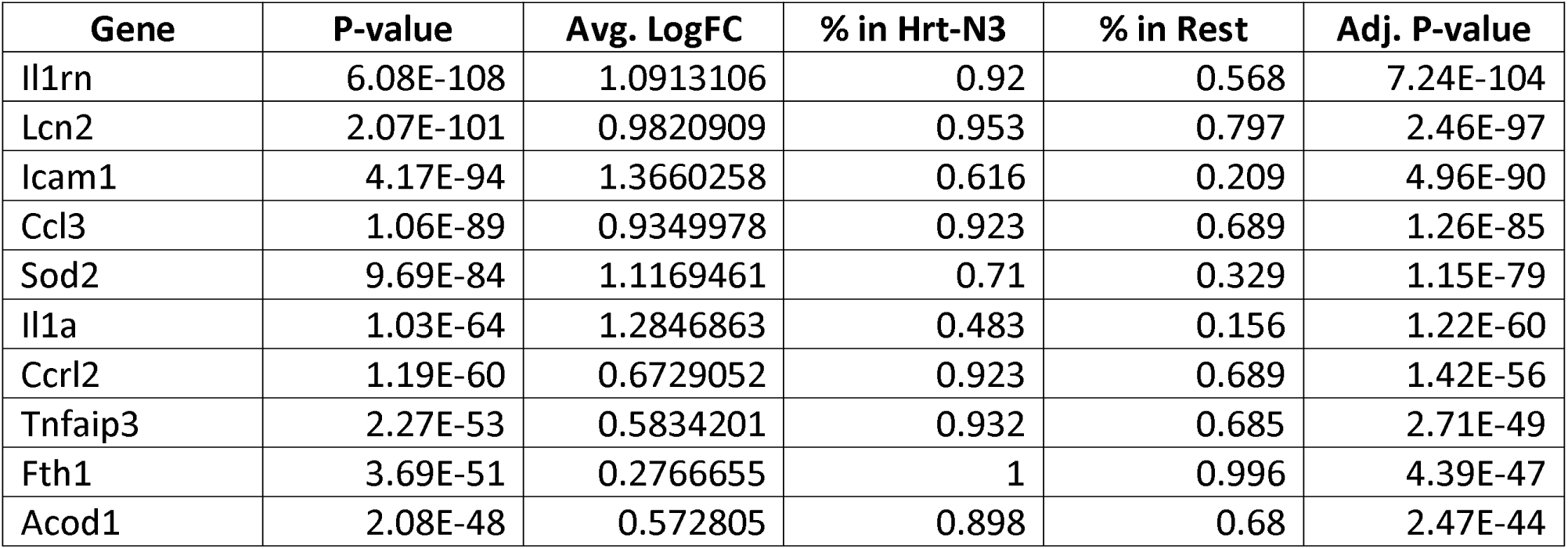
Top 10 differentially expressed genes in intracardiac neutrophil subset Hrt-N3. Wilcoxon Rank Sums test.

**Table 3D.**
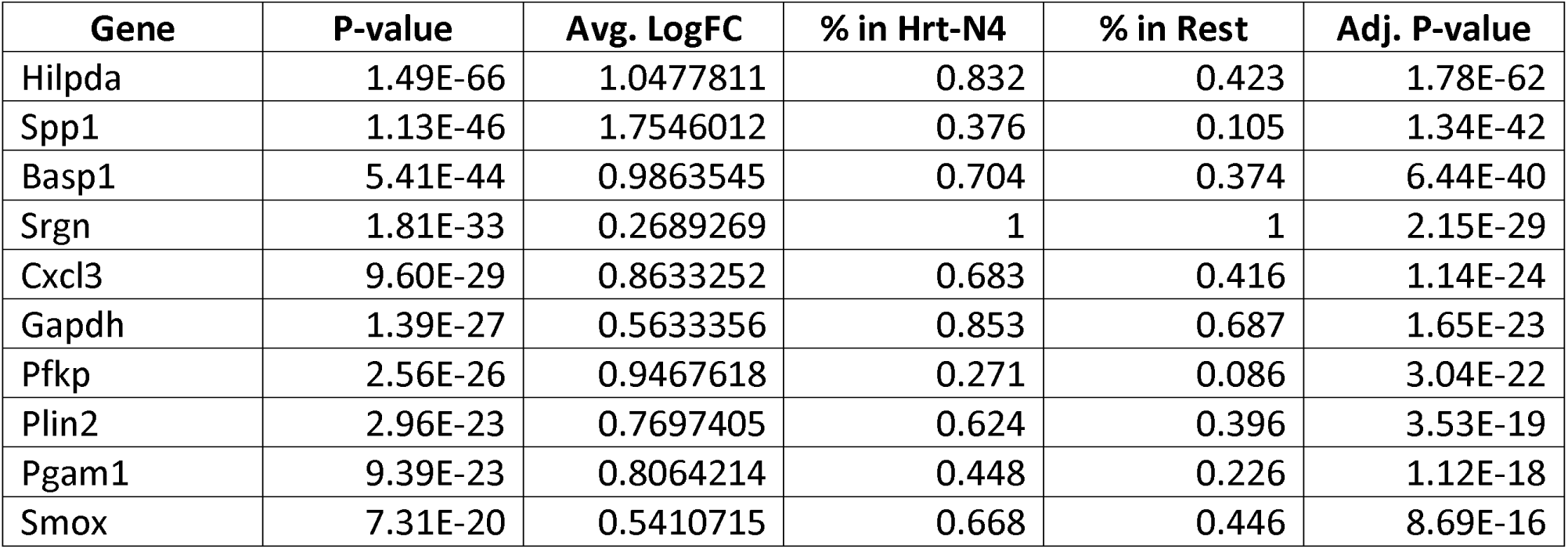
Top 10 differentially expressed genes in intracardiac neutrophil subset Hrt-N4. Wilcoxon Rank Sums test.

**Table 3E.** Top 10 differentially expressed genes in intracardiac neutrophil subset Hrt-N5. Wilcoxon Rank Sums test.

| Gene | P-value | Avg. LogFC | % in Hrt-N5 | % in Rest | Adj. P-value |
| --- | --- | --- | --- | --- | --- |
| Dusp2 | 5.00E-20 | 0.9645395 | 0.595 | 0.342 | 6.19E-16 |
| Tnf | 2.32E-13 | 0.5493787 | 0.795 | 0.663 | 2.87E-09 |
| Gadd45b | 6.45E-12 | 0.4279953 | 0.847 | 0.733 | 7.98E-08 |
| Nfkbia | 8.73E-11 | 0.2569613 | 0.967 | 0.956 | 1.08E-06 |
| Siglecf | 1.92E-10 | 0.7688374 | 0.364 | 0.199 | 2.38E-06 |
| Cst3 | 1.33E-08 | 0.4219797 | 0.71 | 0.629 | 1.64E-04 |
| Hic1 | 6.24E-08 | 0.6748172 | 0.296 | 0.164 | 7.72E-04 |
| Tmem14c | 7.33E-08 | 0.6608851 | 0.31 | 0.182 | 9.08E-04 |
| Eef1a1 | 9.15E-08 | 0.4054992 | 0.751 | 0.693 | 1.13E-03 |
| Dynll1 | 2.99E-07 | 0.4553353 | 0.562 | 0.45 | 3.70E-03 |

Since we and others previously found type I IFN signaling to be detrimental after MI (Cao et al., 2018; King et al., 2017), we inquired whether there were any underlying differences between ISG+ and ISG- neutrophils. Therefore, we bioinformatically isolated and reclustered them. Surprisingly, we recovered all of the same neutrophil subsets in ISG+ as ISG- populations (Figure 2D). To investigate transitions between neutrophil subsets, we performed pseudotime trajectory analysis (Monocle 3.0) on a down-sampled population of cells (n = 5,000 cells) from post-MI day 4 data (Qiu et al., 2017b). Whether analyzed together or individually, ISG+ and ISG- neutrophils resulted in mirrored trajectories (Figure 2E), reinforcing the view that ISG-expression is orthogonal to the other axes of neutrophil differentiation. To examine how ISG expression varies along the neutrophil maturation axis, we compared its expression to *Retnlg* and *Siglecf* (Figure 2F, Figure S5), which served as markers of early and late intracardiac differentiation. We found that recently infiltrated *Retnlg^HI^Siglecf^LOW^* neutrophils expressed the highest levels of ISGs, suggesting that ISG-expression is induced prior to entry into the infarcted heart. Taken together, these findings suggest that MI induces a subset of neutrophils to express ISGs prior to entering the infarcted heart and that ISG expression does not limit intracardiac differentiation potential. In addition, these results provide the first detailed mapping of neutrophil differentiation within the infarcted heart and reveal a previously unrecognized late *SiglecF*+ neutrophil whose function has yet to be explored.

### Monocytes and macrophages within the infarcted heart consist of mirrored ISG+ and ISG- subsets and one exclusively ISG-negative macrophage subset

We next bioinformatically isolated intracardiac monocytes and macrophages and reclustered them (Figure 3A). This revealed Hrt-m1 monocytes (expressing *Ccr2, Plac8,* and *Ms4a4c)*; an ISG+ Hrt-m2 population (Hrt-m2) that appeared to consist of monocytes and macrophages; and the following macrophage subsets – Hrt-M3 (expressing *C1qa C1qb, Mrc1,* and *Ctsd*, including Ccr2- resident macrophages); Hrt-M4 (expressing *Arg1* and *Vegfa*); Hrt-M5 (expressing *Gclm* and *Slc40a1,* which we later reveal to be Nrf2-activated macrophages); and Hrt-M6 (expressing cell cycle genes implying proliferation – not shown) (Figure 3B, Figure S6A-B, Table 4A-E). We termed the ISG+ Hrt-m2 cells monocyte-derived IFN-induced cells (mIFNICs) to distinguish them from neutrophil nIFNICs.

**Figure 3.**
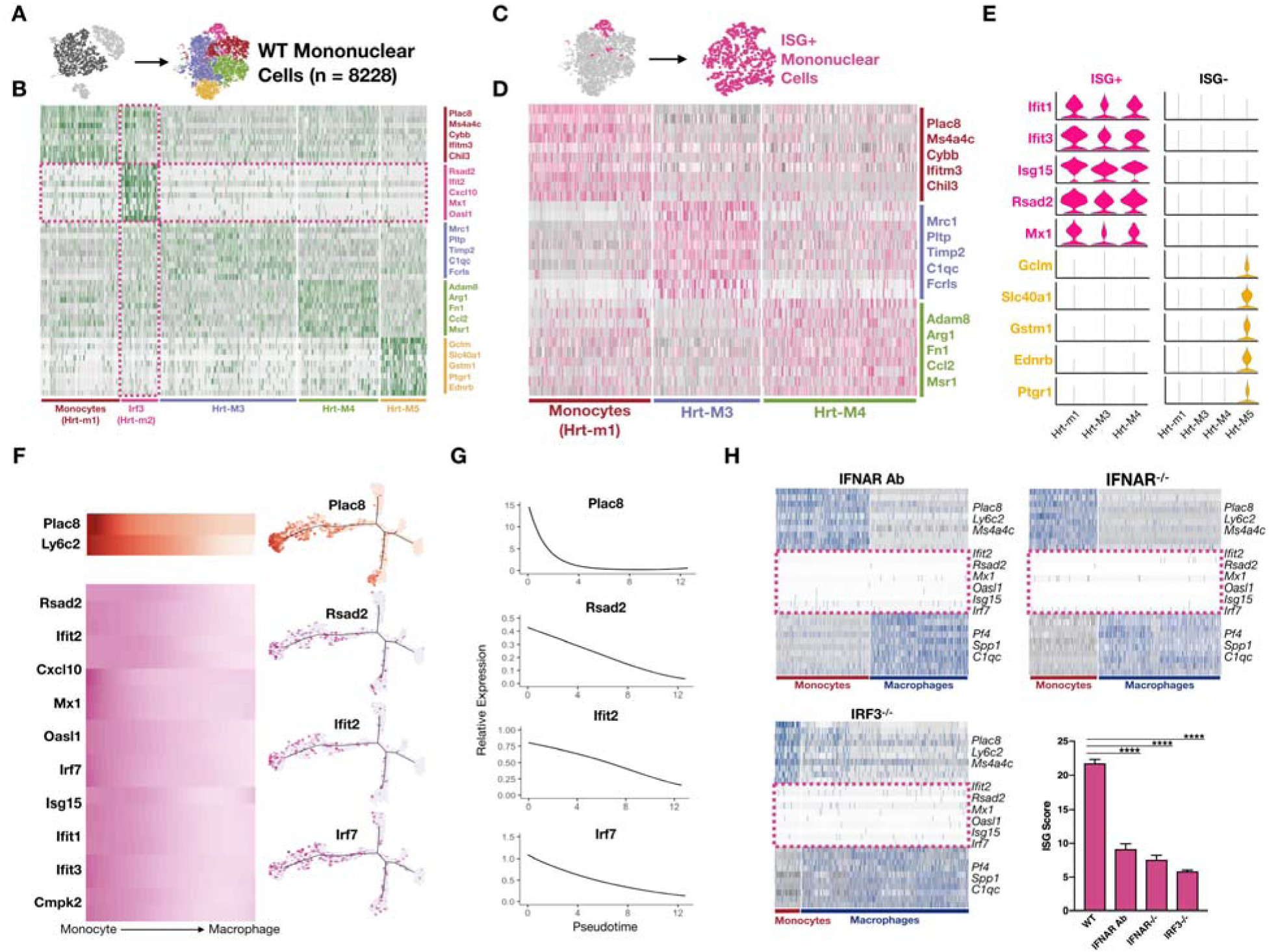
Monocytes and macrophages differentiate into ISG+ and ISG- subsets except for one exclusively ISG- macrophage within the infarcted mouse heart. **(A-H)** Single cell RNA-Seq data of monocytes and macrophages isolated from infarcted cardiac tissue on day 4 after MI. (**A**) t-SNE map illustrating monocytes and macrophages subsetting and reclustering in wildtype (WT) mice. (**B**) Representative heatmap of subset and reclustered monocyes and macrophages (n = 8,228 cells) in WT mice. Dividing cells were bioinformatically removed. (**C,D**) Subclustering of ISG+ mononuclear cells recapitulates all monocyte and macrophage subsets with the exception of Hrt-M5. (**E**). Violin plots of subclustered ISG+ and ISG- cells with ISGs and Hrt-M5 marker genes. (**F,G**) Pseudotime analysis shows ISG expression is highest in Plac8^HI^ Ly6c2^HI^ (monocytes) and declines with differentiation. (**H**) Heatmaps and ISG scores highlighting ISG expression in monocytes and macrophages of IFNAR Ab treated, IFNAR-/-, and IRF3-/- mice. **** P< .0001, Mann-Whitney test.

**Table 4A.** Top 10 differentially expressed genes in intracardiac monocyte subset Hrt-m1. Wilcoxon Rank Sums test.

| Gene | p_val | avg_logFC | pct.1 | pct.2 | p_val_adj |
| --- | --- | --- | --- | --- | --- |
| Ifitm6 | 2.08E-69 | 1.5701412 | 0.566 | 0.073 | 2.67E-65 |
| Hp | 3.47E-66 | 1.5424927 | 0.618 | 0.097 | 4.46E-62 |
| Plac8 | 4.18E-62 | 1.4599853 | 0.993 | 0.449 | 5.37E-58 |
| Itgal | 2.08E-57 | 1.3409417 | 0.544 | 0.084 | 2.67E-53 |
| Sell | 1.15E-42 | 1.25542 | 0.419 | 0.064 | 1.47E-38 |
| Napsa | 3.97E-41 | 1.3395312 | 0.566 | 0.137 | 5.09E-37 |
| Vcan | 5.72E-41 | 1.2499742 | 0.434 | 0.074 | 7.34E-37 |
| Actg1 | 3.37E-32 | 0.6203813 | 0.993 | 0.964 | 4.33E-28 |
| Gsr | 2.82E-30 | 1.1015892 | 0.61 | 0.214 | 3.62E-26 |
| Coro1a | 1.14E-29 | 0.9251717 | 0.846 | 0.497 | 1.47E-25 |

**Table 4B.** Top 10 differentially expressed genes in intracardiac monocyte subset Hrt-m2. Wilcoxon Rank Sums test.

| Gene | p_val | avg_logFC | pct.1 | pct.2 | p_val_adj |
| --- | --- | --- | --- | --- | --- |
| Ifit2 | 5.42E-188 | 2.520278 | 0.84 | 0.037 | 6.96E-184 |
| Ifit3 | 1.24E-163 | 2.4920708 | 0.84 | 0.052 | 1.59E-159 |
| Ifit1bl1 | 1.12E-155 | 2.127182 | 0.555 | 0.009 | 1.43E-151 |
| Rsad2 | 2.71E-150 | 2.9841967 | 0.891 | 0.078 | 3.48E-146 |
| Oasl1 | 2.48E-134 | 2.1394677 | 0.765 | 0.056 | 3.18E-130 |
| Cxcl10 | 2.93E-129 | 2.9161153 | 0.706 | 0.048 | 3.77E-125 |
| Cmpk2 | 3.03E-117 | 1.9633156 | 0.605 | 0.034 | 3.89E-113 |
| Ifit1 | 2.33E-112 | 2.0832972 | 0.639 | 0.044 | 2.99E-108 |
| Mx1 | 5.94E-111 | 1.8898354 | 0.672 | 0.052 | 7.63E-107 |
| Ifit3b | 5.63E-105 | 1.6557469 | 0.437 | 0.013 | 7.23E-101 |

**Table 4C.** Top 10 differentially expressed genes in intracardiac monocyte subset Hrt-M3. Wilcoxon Rank Sums test.

| Gene | p_val | avg_logFC | pct.1 | pct.2 | p_val_adj |
| --- | --- | --- | --- | --- | --- |
| Itm2b | 2.75E-44 | 0.8707329 | 0.979 | 0.919 | 3.53E-40 |
| Pltp | 2.76E-38 | 1.3064994 | 0.678 | 0.223 | 3.54E-34 |
| C1qa | 5.66E-37 | 1.1677891 | 0.853 | 0.437 | 7.27E-33 |
| C1qc | 7.14E-36 | 1.2080658 | 0.86 | 0.502 | 9.17E-32 |
| C1qb | 1.03E-35 | 0.985766 | 0.923 | 0.589 | 1.32E-31 |
| Ccl8 | 1.40E-32 | 1.9525453 | 0.434 | 0.098 | 1.80E-28 |
| ApoE | 4.31E-32 | 1.0312886 | 0.937 | 0.818 | 5.53E-28 |
| Fcrls | 8.60E-30 | 1.1567704 | 0.427 | 0.102 | 1.10E-25 |
| Mrc1 | 4.39E-25 | 0.9100284 | 0.671 | 0.284 | 5.64E-21 |
| Timp2 | 1.34E-22 | 0.8371619 | 0.72 | 0.343 | 1.72E-18 |

**Table 4D.**
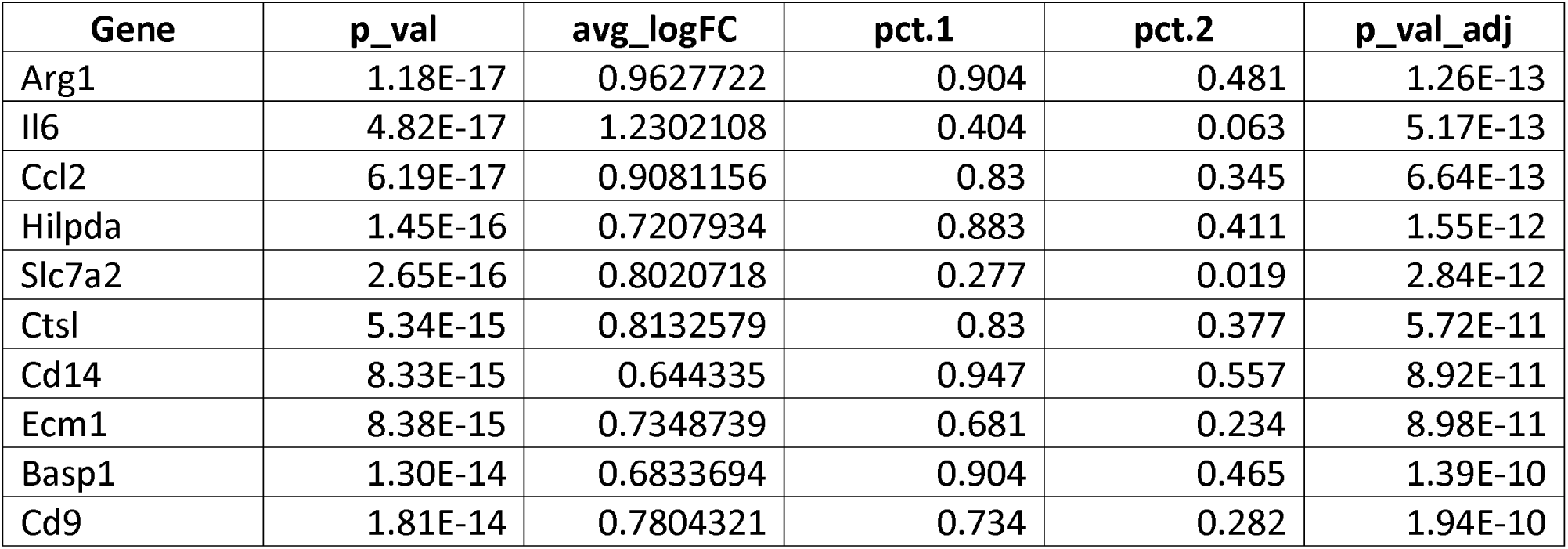
Top 10 differentially expressed genes in intracardiac monocyte subset Hrt-M4. Wilcoxon Rank Sums test.

| Gene | p_val | avg_logFC | pct.1 | pct.2 | p_val_adj |
| --- | --- | --- | --- | --- | --- |
| Arg1 | 1.18E-17 | 0.9627722 | 0.904 | 0.481 | 1.26E-13 |
| Il6 | 4.82E-17 | 1.2302108 | 0.404 | 0.063 | 5.17E-13 |
| Ccl2 | 6.19E-17 | 0.9081156 | 0.83 | 0.345 | 6.64E-13 |
| Hilpda | 1.45E-16 | 0.7207934 | 0.883 | 0.411 | 1.55E-12 |
| Slc7a2 | 2.65E-16 | 0.8020718 | 0.277 | 0.019 | 2.84E-12 |
| Ctsl | 5.34E-15 | 0.8132579 | 0.83 | 0.377 | 5.72E-11 |
| Cd14 | 8.33E-15 | 0.644335 | 0.947 | 0.557 | 8.92E-11 |
| Ecm1 | 8.38E-15 | 0.7348739 | 0.681 | 0.234 | 8.98E-11 |
| BasP1 | 1.30E-14 | 0.6833694 | 0.904 | 0.465 | 1.39E-10 |
| Cd9 | 1.81E-14 | 0.7804321 | 0.734 | 0.282 | 1.94E-10 |

**Table 4E.**
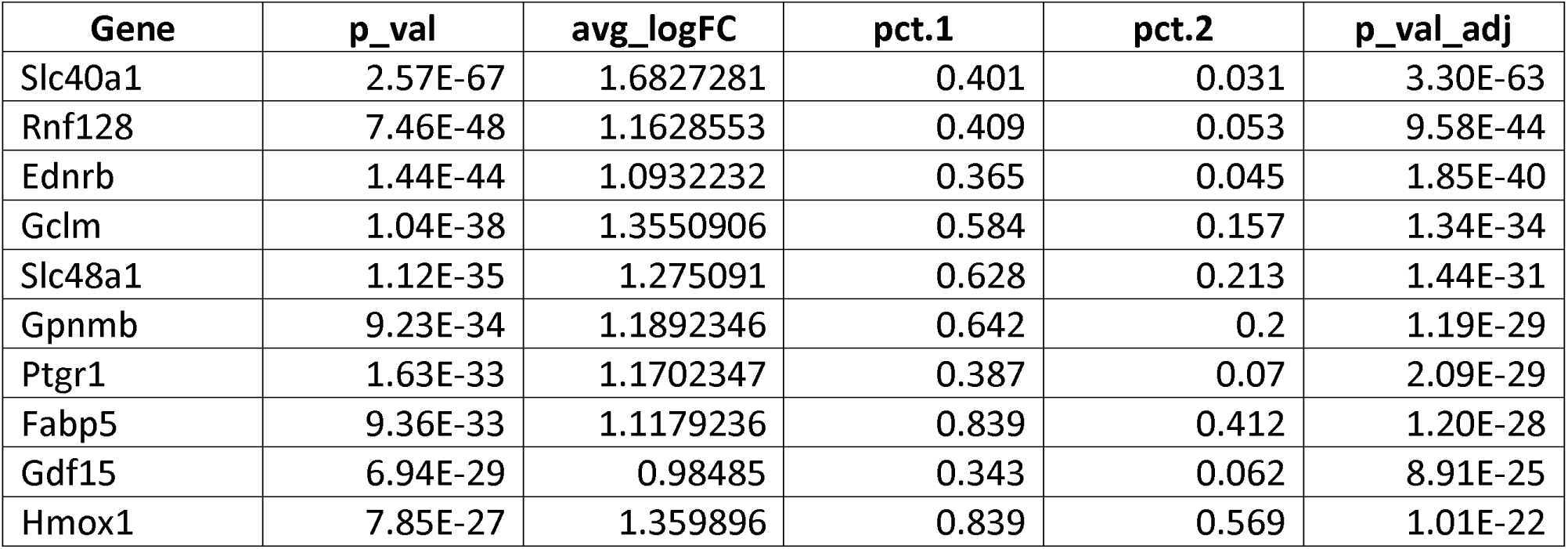
Top 10 differentially expressed genes in intracardiac monocyte subset Hrt-M5. Wilcoxon Rank Sums test.

**Table 4F.**
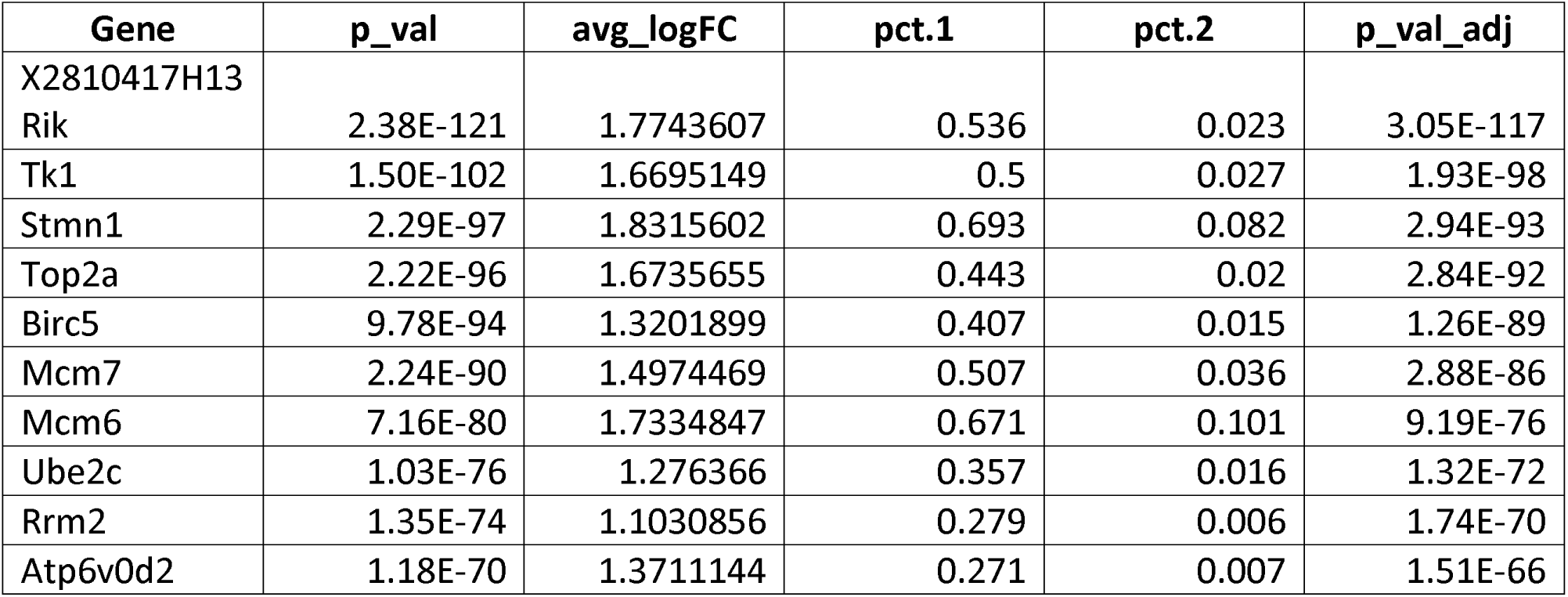
Top 10 differentially expressed genes in intracardiac monocyte subset Hrt-M6. Wilcoxon Rank Sums test.

Similar to our analysis of neutrophils, we bioinformatically isolated ISG+ and ISG- monocytes and macrophages and separately reclustered them (Figure 3C, Figure S6C). This revealed mirrored ISG+ and ISG- subpopulations of each monocyte-macrophage subset except for the ISG-expressing Hrt1-m2 cells, which were exclusively ISG+ by design, and the uncharacterized Hrt-M5 macrophages, which were the only subset to have ISG- cells without a companion ISG+ subpopulation (Figure 3C-E, Figure S6C). Analogous to neutrophils, pseudotime analysis of post-MI day 4 samples showed that ISG expression was highest in recently-infiltrated cells (monocytes and recently differentiated macrophages) expressing *Ly6c2, Plac8,* and *Ms4a4c*, and declined in more mature macrophages regardless of subset (Figure 3F-G, S7). This result, which was confirmed in FACS sorted cells by qPCR (Figure S8A-C) suggested that ISGs mark the earliest monocyte-derived cells of the infarct. ISG expression was almost completely dependent on the transcription factor, interferon regulatory factor 3 (*Irf3),* and the interferon alpha receptor (*Ifnar),* since single cell transcriptomes from the infarcted hearts of *Irf3*^-/-^ mice, *Ifnar*^-/-^ mice, or *WT* mice treated with Ifnar neutralizing antibody showed near complete abrogation of ISG expression (Sheehan et al., 2006) (Figure 3H). Collectively, these data demonstrate that ISGs are induced prior to monocyte arrival to the infarcted heart in an Irf3- and Ifnar-dependent manner. Once in the heart, ISG+ and ISG- monocytes have similar differentiation potentials with the exception of Hrt-M5 macrophages, which had no ISG+ subset. Because of this unique asymmetry, we turned our attention to the Hrt-M5 macrophages.

### Nrf2-dependent signaling in Ccr2- resident macrophages negatively regulates post-MI type I IFN responses

We noticed that the marker genes of Hrt-M5 were genes previously associated with the cytoprotective transcriptional regulator Nrf2 (Nuclear factor (erythroid-derived 2)-like 2) based on chromatin immunoprecipitation experiments or the presence of Nrf2-binding antioxidant response element (ARE) sequences in their promoters (Lee et al., 2003; Malhotra et al., 2010; Moi et al., 1994). To determine whether the Hrt-M5 macrophage population was indeed Nrf2-dependent, we subjected *Nrf2^-/-^* mice to MI and examined the resulting single cell transcriptomes of the mononuclear cells (Figure 4A). We observed dramatic reductions of the Hrt-M5 marker genes but recovered all of the other ISG+ and ISG- intracardiac mononuclear populations defined above (Figure 4B). Volcano plots comparing *Nrf2^-/-^* and *WT* macrophage subsets further demonstrated that reductions in Nrf2-dependent genes did not substantially alter the underlying cell type (Figure 4C). Based on the abrogation of its marker genes, we concluded that Hrt-M5 cells are Nrf2-activated macrophages and we termed its marker genes Nrf2-stimulated genes (NSGs). Using the top NSGs, we then created a composite NSG score (analogous to the ISG score above) and confirmed it was significantly reduced in Nrf2-/- mice after MI compared to WT (Figure 4D).

**Figure 4.**
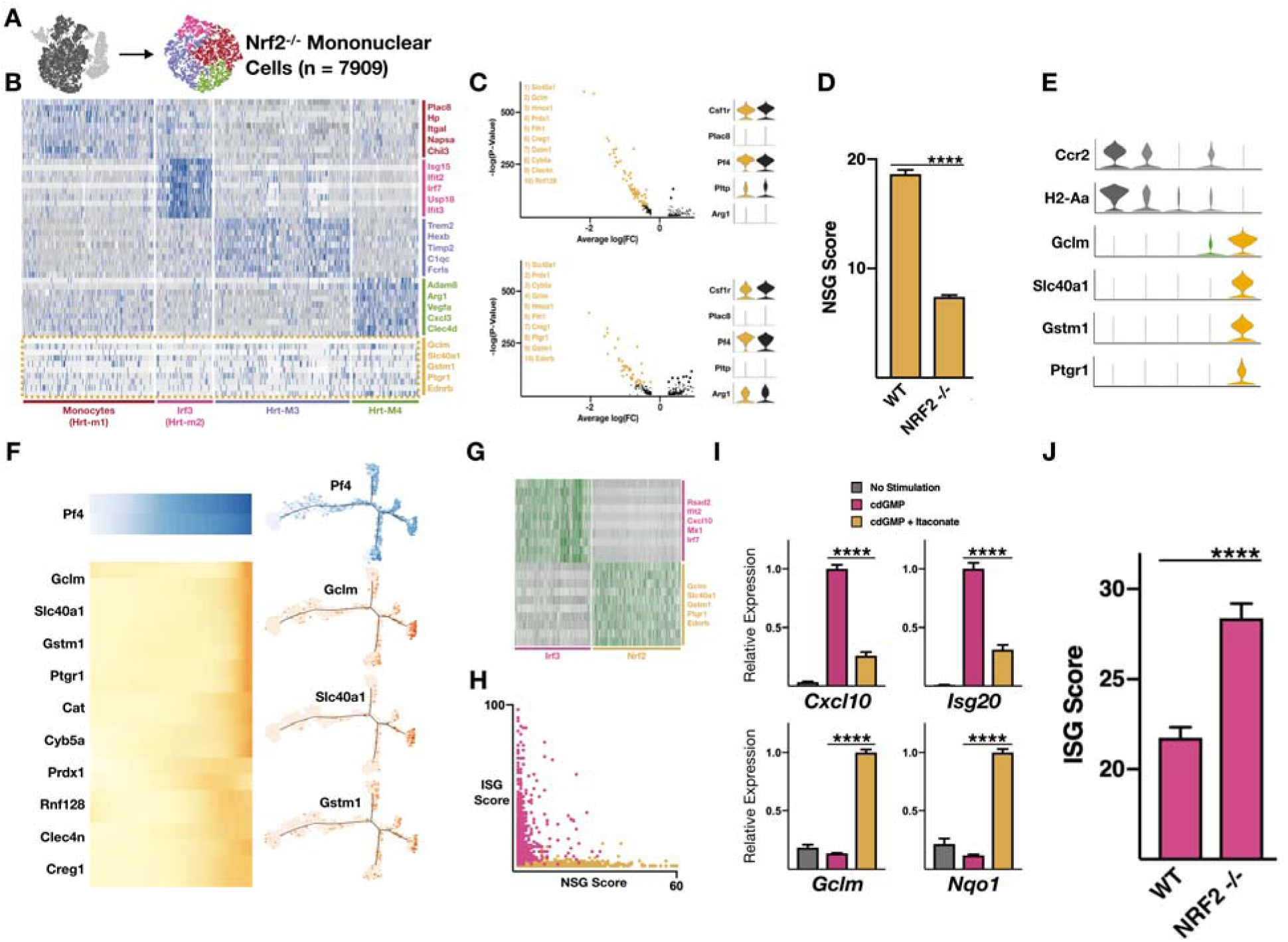
*Ccr2*- resident macrophages impose Nrf2-dependent inhibition on ISG expressing bone marrow-derived myeloid cells in the infarcted heart. (**A-D**) Single cell RNA-Seq data from *Nrf2-/-* mice on day 4 after MI. (**A**) t-SNE map of monocytes and macrophages subset and reclustered. (**B**) Heatmap of subset and reclustered mono-macs from *Nrf2-/-* infarcted hearts. Gold line highlights Nrf2-stimulated genes (NSGs). (**C**) Down-regulated genes in Nrf2-/- compared to WT macrophages (Mac01 and Mac02). (**D**) Nrf2-stimulated gene (NSG) score (defined as the summed expression of top Nrf2-stimulatyed genes, see methods) in WT and Nrf2 -/- monocytes and macrophages. (**E**) *CCR2* and *H2-Aa* expression of NSG+ cells (gold). (**F**) Pseudotime analysis highlighting NSG expression in macrophages. (**G**) Heatmap illustrating dichotomous expression of ISGs and NSGs in wild type mice on day 4 after MI. (**H**) Scatterplot of ISG-score and NSG-score for WT mono-mac single cells. **G**) qPCR of ISGs (*Cxcl10, Isg20*), and NSGs (*Gclm, Nqo1*) from bone marrow derived macrophages treated with cyclic-di-GMP and itaconate. (**J**) ISG scores of monocytes and macrophages from *WT* and *Nrf2 -/-* mice D4 after MI. **** P< .0001, Mann-Whitney test.

Comparisons between the expression of proinflammatory ISGs and cytoprotective NSGs revealed a curious dichotomy – ISGs and NSGs were almost never induced in the same cell (Figure 4E). Plotting the ISG and NSG scores of single mononuclear cells against each other confirmed the mutually exclusive expression of Irf3- and Nrf2-dependent gene expression programs (Figure 4F). To explain the dichotomy, we considered the possibility that Nrf2-activated cells are resident macrophages. We reasoned that ISGs are induced remotely (outside of the heart) because they were found in peripheral blood leukocytes of mice and humans. This would leave resident macrophages as the only myeloid cells in the infarct to have “never seen” remote ISG- inducing stimuli. Indeed, NSG-expressing Hrt-M5 macrophages were the only subset not to have an ISG-expressing subpopulation. In addition, they were Ccr2-, which is a unique attribute of resident macrophages that discriminates them from recently recruited bone-marrow derived macrophages (Figure 4G). Also supporting this model was pseudotime trajectory analysis, which showed that NSGs were most lowly expressed by monocytes and most highly expressed by the most mature and differentiated cardiac macrophages, consistant with their residing in the heart prior to infarct (Figure 4H). Taken together, this suggests that Nrf2-activated Hrt-M5 cells are resident macrophages and that the ISG-NSG dichotomy is explained by differences in macrophage tissue origin.

Nrf2 negatively regulates inflammation in many models through incompletely-defined mechanisms (Ahmed et al., 2017; Lee et al., 2005; Mills et al., 2018; Thimmulappa et al., 2006; Yu et al., 2019). To test whether Nrf2 signaling negatively inhibits expression of ISGs, we performed mechanistic experiments in vitro and in vivo. Bone marrow derived macrophages were stimulated with the Irf3-inducing cyclic dinucleotide, cyclic-di-GMP (cdGMP), with and without exogenous Nrf2-activating 4-O-itaconate (4OI) (Mills et al., 2018). Whereas cdGMP alone increased the expression of several ISGs, co-stimulation with cdGMP and 4OI significantly decreased the expression of ISGs while increasing the expression of NSGs (Figure 4I). *In vivo,* we observed significantly increased levels of ISGs in infarcted Nrf2-deficient mice compared to *WT* mice as quantified by the ISG score (Figure 4J), suggesting the presence of Nrf2-dependent inhibition of ISG expression. Taken together, these results suggest that Nrf2-activation in Ccr2- resident macrophages negatively regulates pathologic ISG expression in infiltrating bone marrow-derived monocytes and neutrophils after MI.

### ISG expression neutrophil and monocyte subsets in the bone marrow after MI

Since ISG expression was identified in multiple myeloid lineages (neutrophils and monocytes) of the heart and blood, we hypothesized that ISG activation may be initiated within the bone marrow, where myeloid cells arise from a common progenitor. We performed single cell RNA-Seq on bone marrow hematopoietic cells on days 1-3 after MI and found ISG expression in subsets of developing neutrophils and monocytes. Cluster analysis identified immature neutrophils (BM-N1) expressing *Camp* and *Chil3,* a spontaneously clustering ISG+ population (BM-N2), and mature neutrophils (BM-N3) expressing *Ccl6* and *Clec4d* (Figure 5A-B). We bioinformatically isolated ISG+ and ISG- cells, separately reclustered them, and recovered ISG+ neutrophils spanning the spectrum of immature and mature neutrophils (Figure 5C). Pseudotime analysis confirmed this finding by defining a monotonic progression from immature BM-N1 (*Camp^HI^*) to mature BM-N3 (*Ccl6^HI^*) neutrophils (Figure 5D) and demonstrating that ISG expression increased with neutrophil maturation (Figure 5E). Finally, we arranged and plotted neutrophil transcriptomes from all three tissue compartments, which revealed MI-induced ISG-expressing subsets gradually increase in ISG expression magnitude from bone marrow to blood to heart (Figure S9).

**Figure 5.**
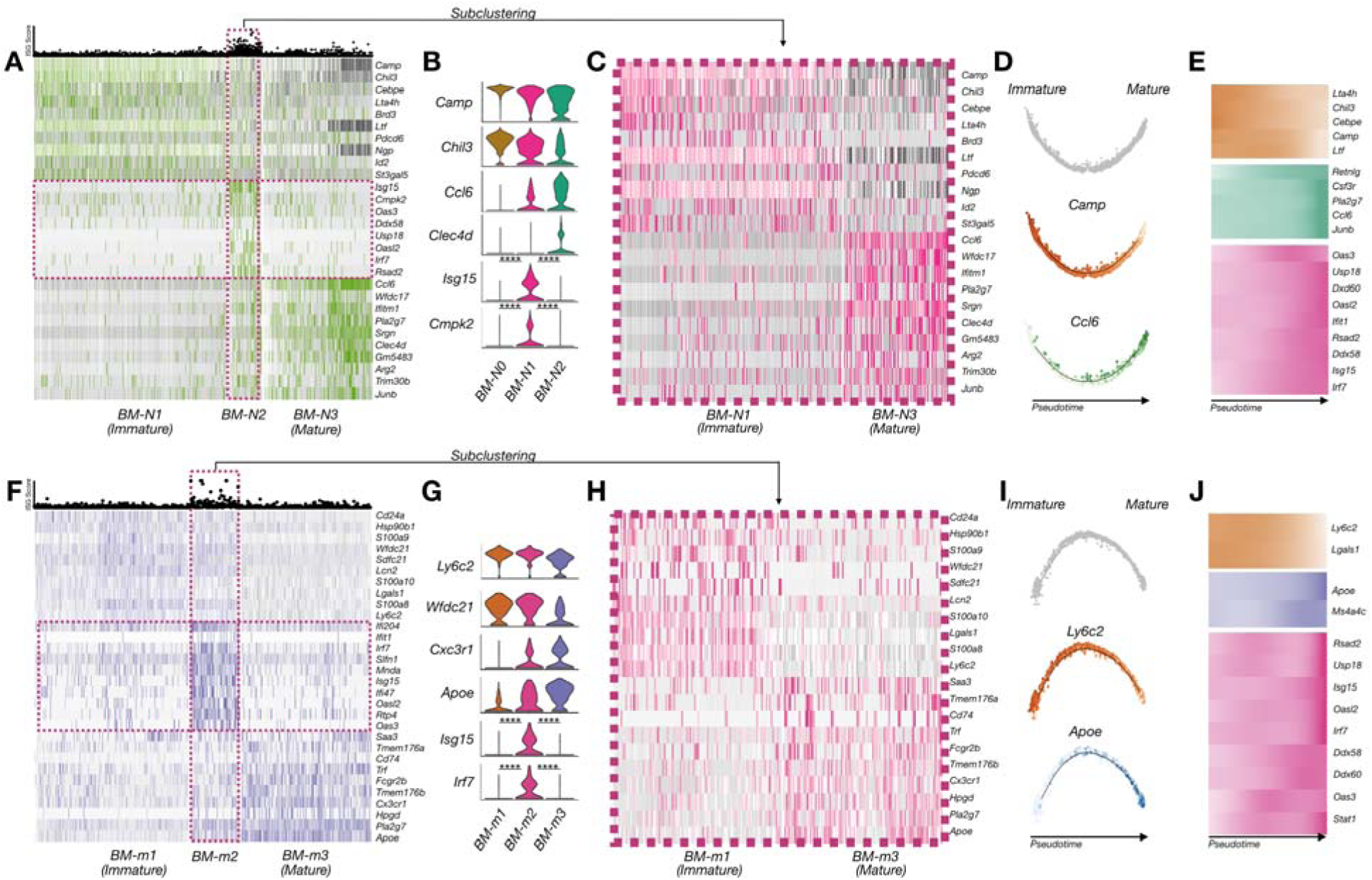
MI induces mirrored ISG+ and ISG- neutrophil and monocyte subsets in the bone marrow. Single cell RNA-Seq data of neutrophils (A-E) and monocytes (F-J) isolated from bone marrow of WT mice on D1-D3 after MI. (**A,F**) Heatmap and ISG scores of bone marrow (**A**) neutrophils (n = 4473 cells, 3 mice) and (**F**) monocytes (n = 1382 cells, 3 mice) of combined data from days 1, 2 and 3 after MI arranged from immature to mature. ISG+ group highlighted in pink. (**B,G**) Violin plots showing maturation markers and ISGs in (**B**) neutrophils and monocytes (**G**). (**C,H**) Heatmap of isolated and reclustered ISG+ (**C**) neutrophils and (**H**) monocytes reveals immature and mature subsets. (**D,E,I,J**) Pseudotime trajectory map of (**D**) neutrophil and (**I**) monocyte maturation and heatmap of ISG expression has a function of pseudotime (**E,J**).

We performed a similar analysis on bone marrow monocytes. After subsetting and reclustering, we observed immature monocytes (BM-m1) expressing *Ly6c2* and *Lgals1*, ISG-expressing monocytes (BM-m2), and mature monocytes (BM-m3) expressing *Cx3cr1* and *Apoe* (Figure 5F-G). The ISG+ monocytes could be subset and reclustered to recover the underlying ISG+ immature and ISG+ mature monocytes (Figure 5H), which again demonstrated that activation of the IFN transcriptional program is orthogonal to maturation and does not limit or bias cellular differentiation potential. Pseudotime analysis confirmed the monotonic progression from immature BM-m1 to mature BM-m3 monocytes (Figure 5I), the latter of which closely matched circulating cells in the blood. As anticipated, ISG expression increased as bone marrow monocytes matured (Figure 5J). Similar to neutrophils, we arranged and plotted monocyte transcriptomes from all three compartments, which revealed that MI-induced ISG- expressing subsets gradually increase in ISG expression magnitude from bone marrow to blood to heart (Figure S10). Taken together, these results suggest that ISG expression begins at or before the level of granulocyte-macrophage progenitors (GMPs). Having demonstrated that MI-induced type I IFN signaling begins within the bone marrow, far from the site of tissue injury, we next examined a hyperinflammatory bone marrow pathology to determine whether it increased myeloid type I IFN signaling.

### Tet2-/- mice exhibit spontaneous and heightened ISG expression in bone marrow myeloid progenitors and their neutrophil and monocyte progeny

Clonal hematopoiesis is an age-associated condition leading to myeloid bias, hyperinflammatory phenotypes, increased risk of cardiovascular events, and increased mortality in humans (Jaiswal et al., 2014; Jaiswal et al., 2017). Tet2 (Tet methylcytosine dioxygenase 2), an epigenetic modifier with broad effects on transcription, is one of the most common loss of function mutations associated with clonal hematopoiesis in humans. Mechanistic studies of Tet2-mutated patients and Tet2-deficient mice have taken a candidate approach and focused on monocyte-derived macrophages, IL6, IL1β, and the inflammasome (Bick et al., 2019; Fuster et al., 2017; Jaiswal et al., 2017); however, the importance of other cell types (e.g. neutrophils) and innate immune pathways (e.g. type I IFNs) remains unexplored. We hypothesized that Tet2-deficient mice might have abnormal type I IFN innate immune activation in monocytes and neutrophils of the bone marrow. To test this, we performed single cell RNA-Seq of myeloid cells in the bone marrow of Tet2^-/-^ mice at steady-state and after MI. Compared to WT mice, we observed heightened and spontaneous induction of ISG expression and the ISG score in myeloid progenitors and their neutrophil and monocyte progeny, both at steady state and after MI (Figure 6A-C).

**Figure 6.**
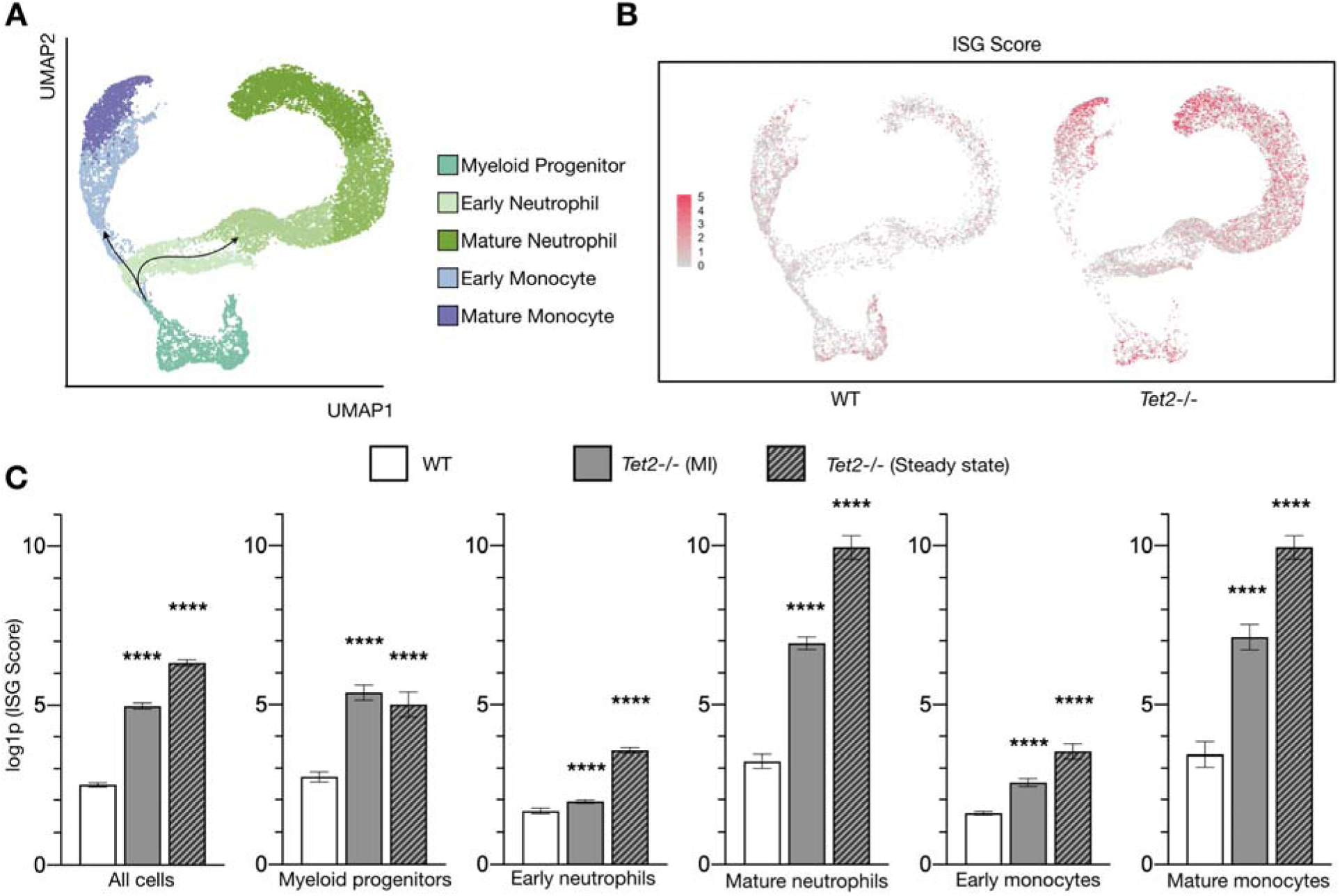
Tet2-deficient mice exhibit potent and spontaneous ISG induction in bone marrow myeloid progenitors and their progeny at steady state and after MI. **(A-C)** Single-cell RNA-seq data of bone marrow myeloid cells from WT control mouse after myocardial infarction (n=5,651 cells) and *Tet2^-/-^* mice in steady-state (n=10,337 cells) and after myocardial infarction (n=13,531 cells). **(A)** Integrated UMAP of all CD11b^+^ bone marrow myeloid cells enriched for Lin^-^Sca1^+^c-Kit^+^ cells. Arrows indicate inferred maturation trajectory along monocyte and neutrophil lineages. **(B)** Feature plot of ISG score in WT control and *Tet2^-/-^* mice after myocardial infarction. **(C)** ISG scores of bone marrow myeloid cells (split by cell type) from WT control mouse after MI and *Tet2^-/-^* mice in steady-state and after MI. Data are displayed as mean ± SEM. **** p<0.0001, Mann-Whitney test.

## Discussion

In summary, our data demonstrate that ischemic injury in the heart induces type I IFN signaling in the bone marrow, far from the site of tissue injury, at the origins of emergency myelopoiesis. We show that this response occurs not only in monocytes and monocyte-derived macrophages but also in neutrophils. We show that the type I IFN response is enhanced in the bone marrow of Tet2-deficient mice, is negatively regulated by Nrf2-dependent signaling originating from Ccr2- resident cardiac macrophages and is Irf3- and Ifnar-dependent. Our findings have translational significance because the type I IFN response, which was previously thought to be inaccessible within the heart, is now readily quantifiable (by calculating the ISG score) in clinically accessible peripheral blood neutrophils and monocytes of patients presenting with acute MI.

Single cell transcriptomics was essential for discrimination of ISG+ and ISG- myeloid subpopulations in our study because sensitive and specific antibodies for flow cytometric analysis of this pathway are lacking and because ensemble measurement techniques obscure the relative contributions of individual myeloid cell subsets (Shalek et al., 2013; Shalek et al., 2014). This functional heterogeneity has pathologic significance because ISGs include many proinflammatory cytokine and chemokine genes (Muller et al., 1994; Shalek et al., 2014) and because we previously showed that genetic or pharmacologic inhibition of IFN signaling reduces post-MI inflammation and protects against adverse ventricular remodeling and death due to ventricular rupture (King et al., 2017).

Our results raise several mechanistic questions for future studies. How does MI initiate IFN signaling in the distant bone marrow and why does it affect only a defined subset of maturing myeloid cells? Previous studies have shown that soluble stimuli such as exogenous poly(I:C) (Essers et al., 2009; Muller et al., 1994) or IFNα (Essers et al., 2009), endogenous IL1β (Sager et al., 2015) can induce proliferation of bone marrow stem and progenitor cells, but they were not shown to do so heterogeneously. It remains unclear how systemic and diffusible substances would activate some but not all neutrophils and monocytes. Sympathetic nervous system discharge and adrenergic signaling would provide a spatiotemporally localized mechanism for induction as would proximity to vasculature or another bone marrow niche, but none of these have been associated with induction of type I IFN signaling or ISG expression (Dutta et al., 2012; Katayama et al., 2006). We and others observed that ISG expression in the heart was dramatically decreased in *cGas^-/-^* mice after MI suggesting that cytosolic DNA sensing may play an important role (Cao et al., 2018; King et al., 2017). Irf3-activating signals downstream of cGAS-dependent DNA sensing are gap junction permeable and may mediate contact-dependent intercellular communication (Ablasser et al., 2013; Patel et al., 2009) with cells of the bone marrow niche (Pinho and Frenette, 2019; Tikhonova et al., 2019). Alternatively, cell-autonomous Irf3 activation resulting from DNA damage (Li and Chen, 2018), micronuclei (Harding et al., 2017; Mackenzie et al., 2017), or transient nuclear rupture (Denais et al., 2016; Raab et al., 2016) could generate heterogeneous populations of IFN-induced cells. Finally, further exploration of the spontaneous activation of type I IFN signaling in the bone marrow of Tet2-deficient mice may provide important mechanistic clues. Our findings also suggest the need to investigate whether type I IFN signaling contributes to the pathologic consequences of Tet2-mutation-associated clonal hematopoiesis in humans.

From a clinical perspective, our data provide the first direct evidence that type I IFN signaling is activated by MI in humans. The ability to quantify ISG expression in peripheral blood leukocyte subsets using single cell transcriptomics will enable correlation with clinical outcomes and response to therapies. Using the ISG score as a biomarker, one can now test whether patients with an excessive type I IFN response to MI are likely to have worse clinical outcomes in terms of arrhythmias, heart failure, recurrent MI, and overall survival. It will be interesting to compare steady state ISG scores with established inflammatory biomarkers (e.g. C reactive protein) and cardiovascular risk (Ridker, 2004; Ridker et al., 2000). Recent studies have demonstrated that anti-cytokine therapy targeting IL1β can reduce recurrent cardiovascular events in humans, and it has been suggested that preferential benefit is seen by subsets of patients in whom excessive inflammation drives the pathology (Ridker et al., 2017). Whether quantification of ISG score can identify patients who may preferentially benefit from anti-cytokine or immune modulatory therapy can now be tested. Finally, if ISG scores predict worse outcomes in patients, the causal role of ISGs can be tested using anti-IFNAR antibody therapy, which has shown acceptable safety profiles during chronic administration in phase 3 clinical trials for lupus (Furie et al., 2017).

## Acknowledgements

We thank the Single Cell Core at Harvard Medical School and the Institute of Genomic Medicine at UC San Diego for technical assistance. The work was funded by AHA17IRG33410543(K.R.K), NIH R00HL129168 (K.R.K.), and DP2AR075321 (K.R.K.); AHA14FTF20380185 (A.D.A.); 5R01HL122208 (R.W.).

## Author Contributions

All authors designed and performed the experiments and analyzed the data. D.M.C., R.P.N.Jr., and K.R.K. wrote the manuscript. All authors reviewed results and revised the manuscript.

## Supplemental Figures

**Figure S1.**
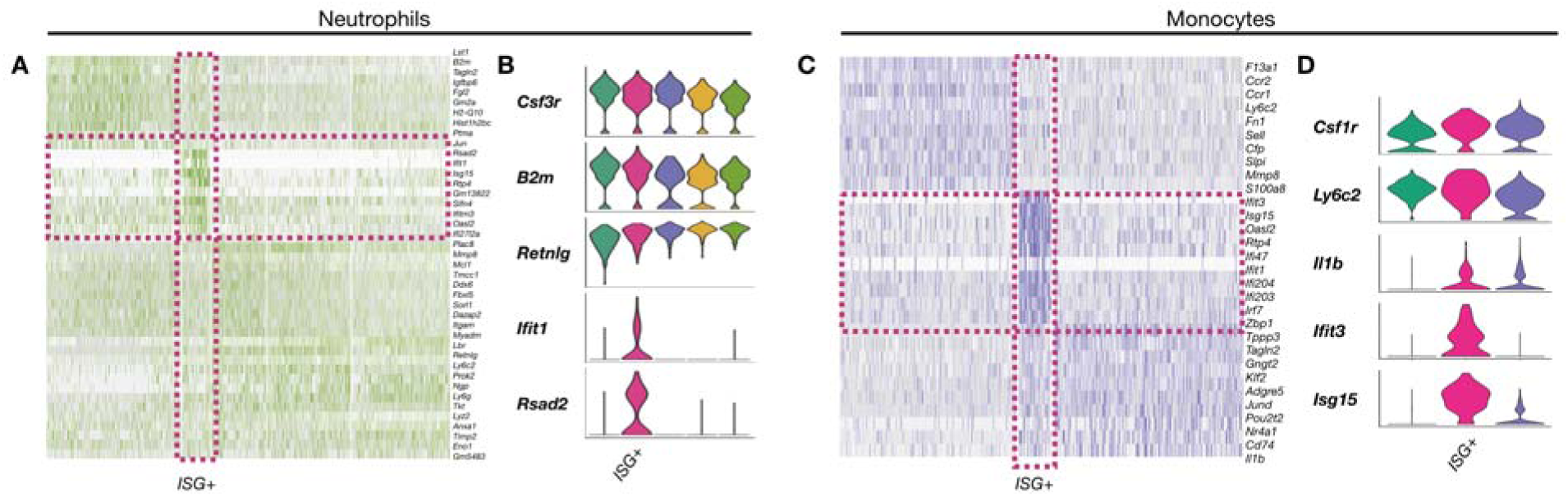
A subset of peripheral blood neutrophils and monocytes express IFN-stimulated genes after MI in mice. **(A-D)** Representative single cell RNA seq data from murine peripheral blood neutrophils and monocytes collected on D2 post-MI. (**A,C**) Representative heatmaps and (**B,D**) violin plot of neutrophils (**A,B**) and monocytes (**C,D**).

**Figure S2.**
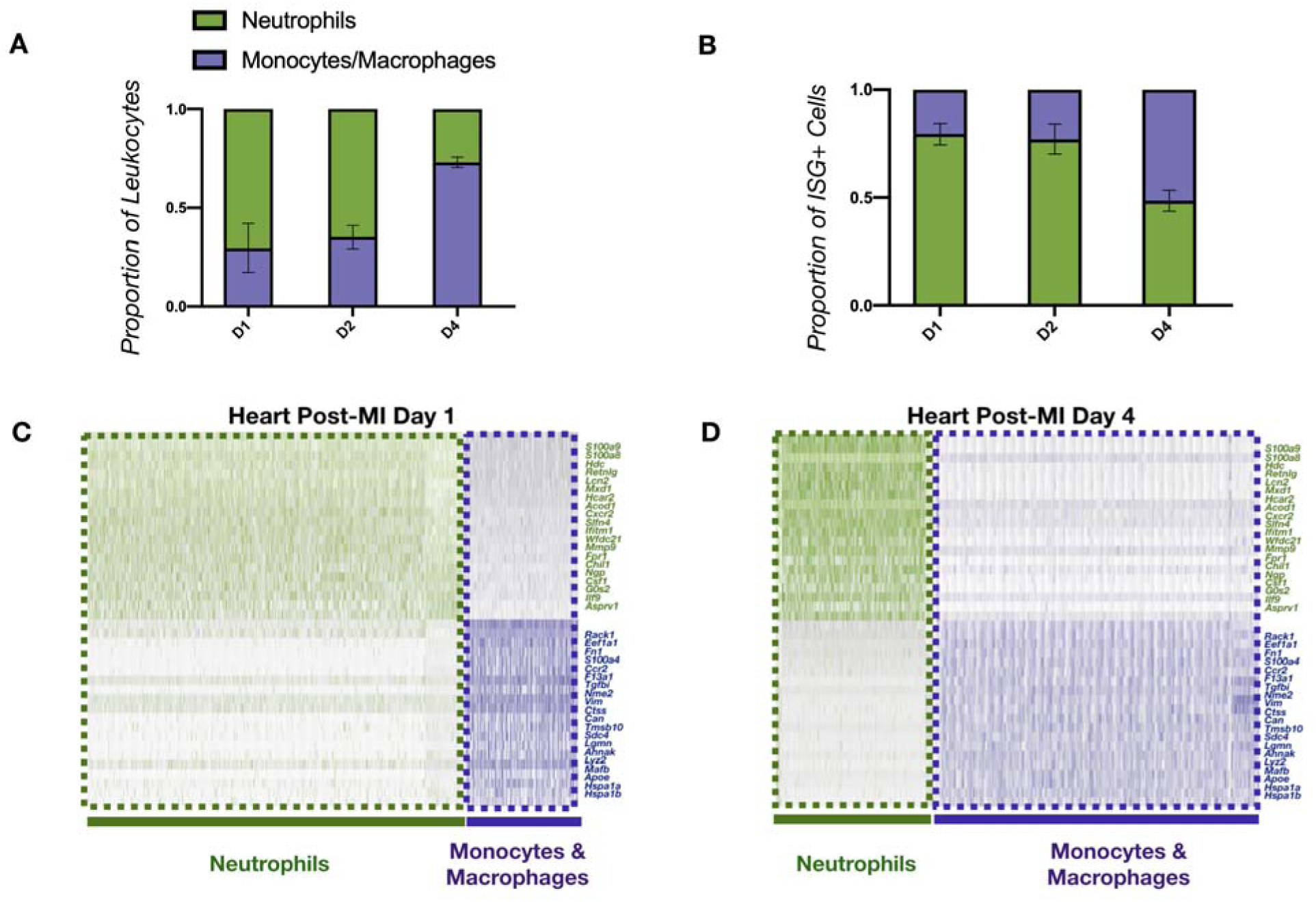
Characterization of MI-induced leukocytosis by single cell RNA seq. (**A**) Relative proportions of neutrophils and monocytes/macrophages as a function of time post-MI (n = 3 mice per day). (**B**) Proportion of ISG+ cells follow a similar trend to bulk migration and cardiac migration. (**C,D**). Representative heatmaps showing neutrophil (green) and monocyte/macrophage (blue) marker genes on (**C**) D1 and (**D**) D4 post-MI.

**Figure S3.**
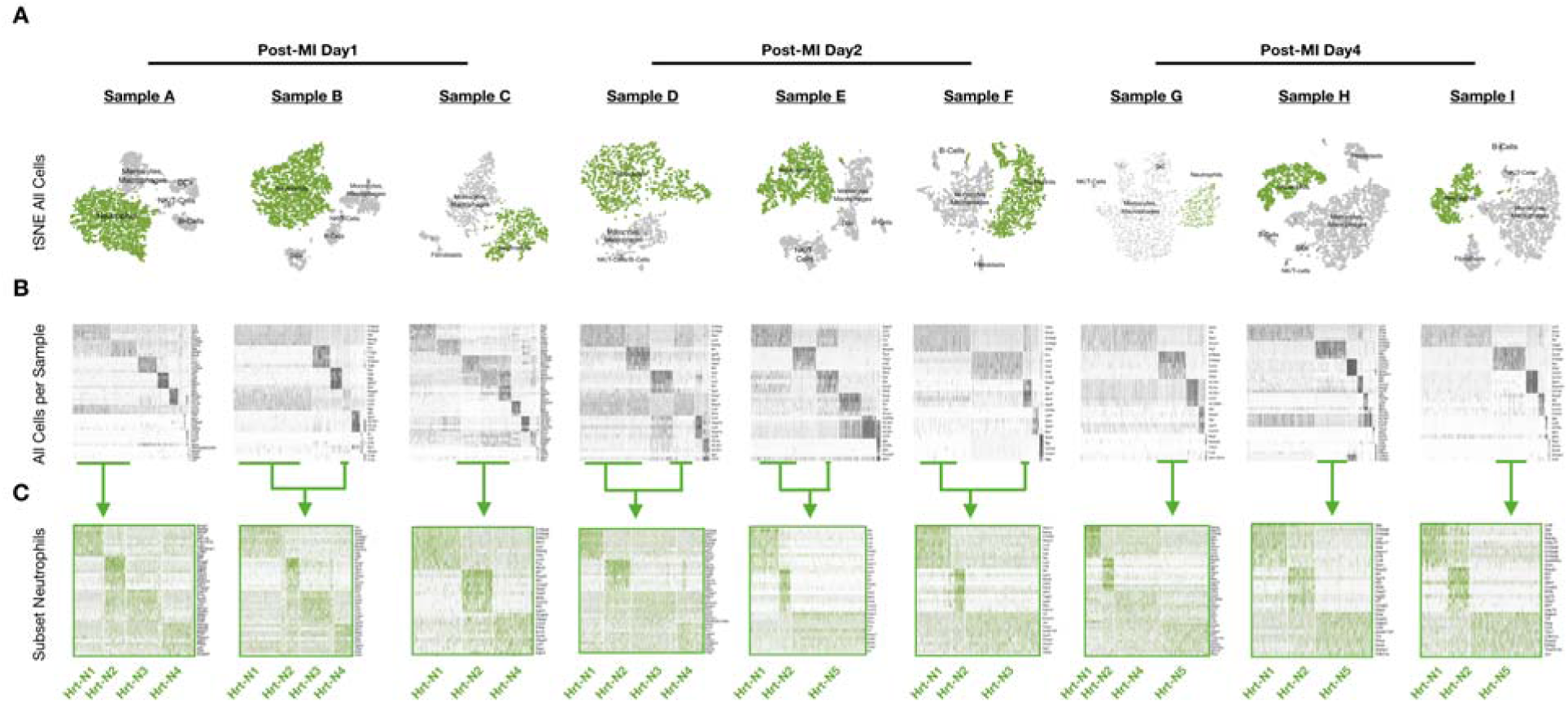
Temporal characterization of mouse intracardiac neutrophil subsets after MI by single-cell RNA seq. (**A**) t-SNE of cells from mouse hearts on days 1-4 after MI. Neutrophils are colored green. (**B**) Clustering of all cells (shown in grey) from each sample uncover neutrophil clusters that can be (**C**) subset, reclustered, and displayed as a heatmap to reveal day-specific neutrophils subsets after MI. Heatmaps (**B,C**) show gene z-score ranging from −2.5 (white), 0 (grey), and +2.5 (dark grey for **B**, green for **C**).

**Figure S4.**
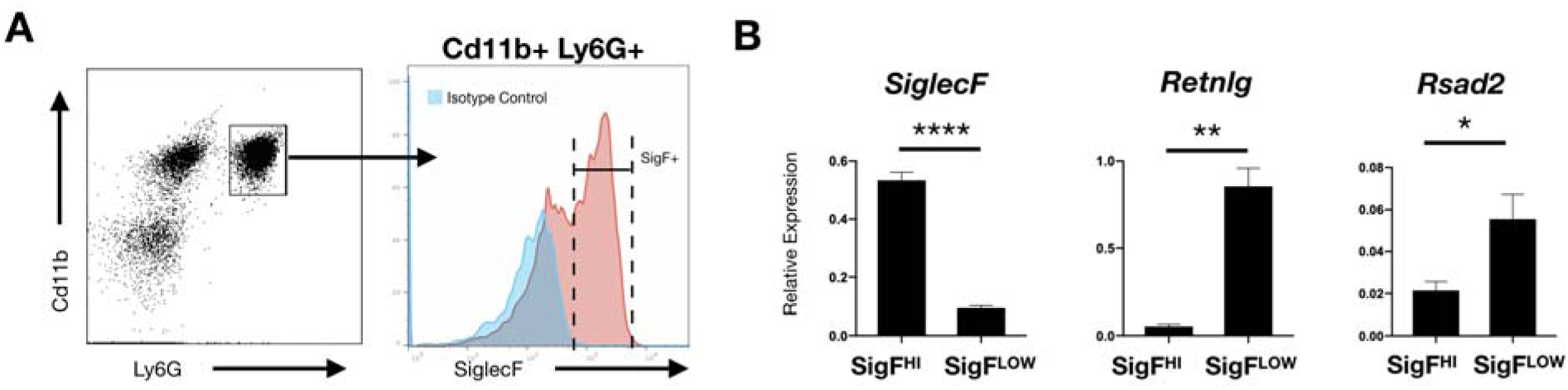
A subset of post-MI cardiac neutrophils express SiglecF. (A) FACS sorting strategy of neutrophils from D4 post permanent LAC ligation of enzymatically digested infarct. (B) qPCR results of SiglecF HI v LOW (n = 6, * *P <.*05, ** *P <.*01,**** *P <.*0001, student t-test).

**Figure S5.**
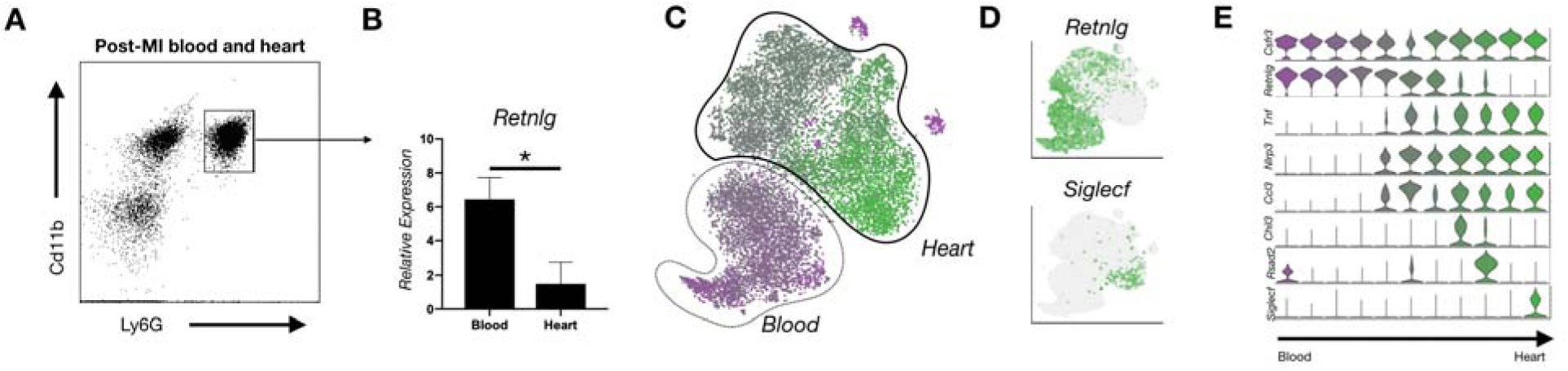
Murine peripheral blood neutrophils are *Retnlg* HI relative to post-MI intracardiac neutrophils by scRNA-seq and qPCR. (**A**) Flow cytometry gating strategy to sort neutrophils from post-MI blood and heart (Cd11b+Ly6G+). (**B**) Bar plot of *Retnlg* expression in neutrophils from blood vs heart. (**C**) t-SNE map of blood (purple) and heart (green) neutrophils (delineated by dotted outlines). (**D**) Expression of *Retnlg* (top) and *Siglecf* (bottom) on t-SNE map. (**E**) Violin plot of neutrophil subsets from blood and heart. Data shown are mean +/- s.e.m. * *P < 0.05,* paired student t-test.

**Figure S6.**
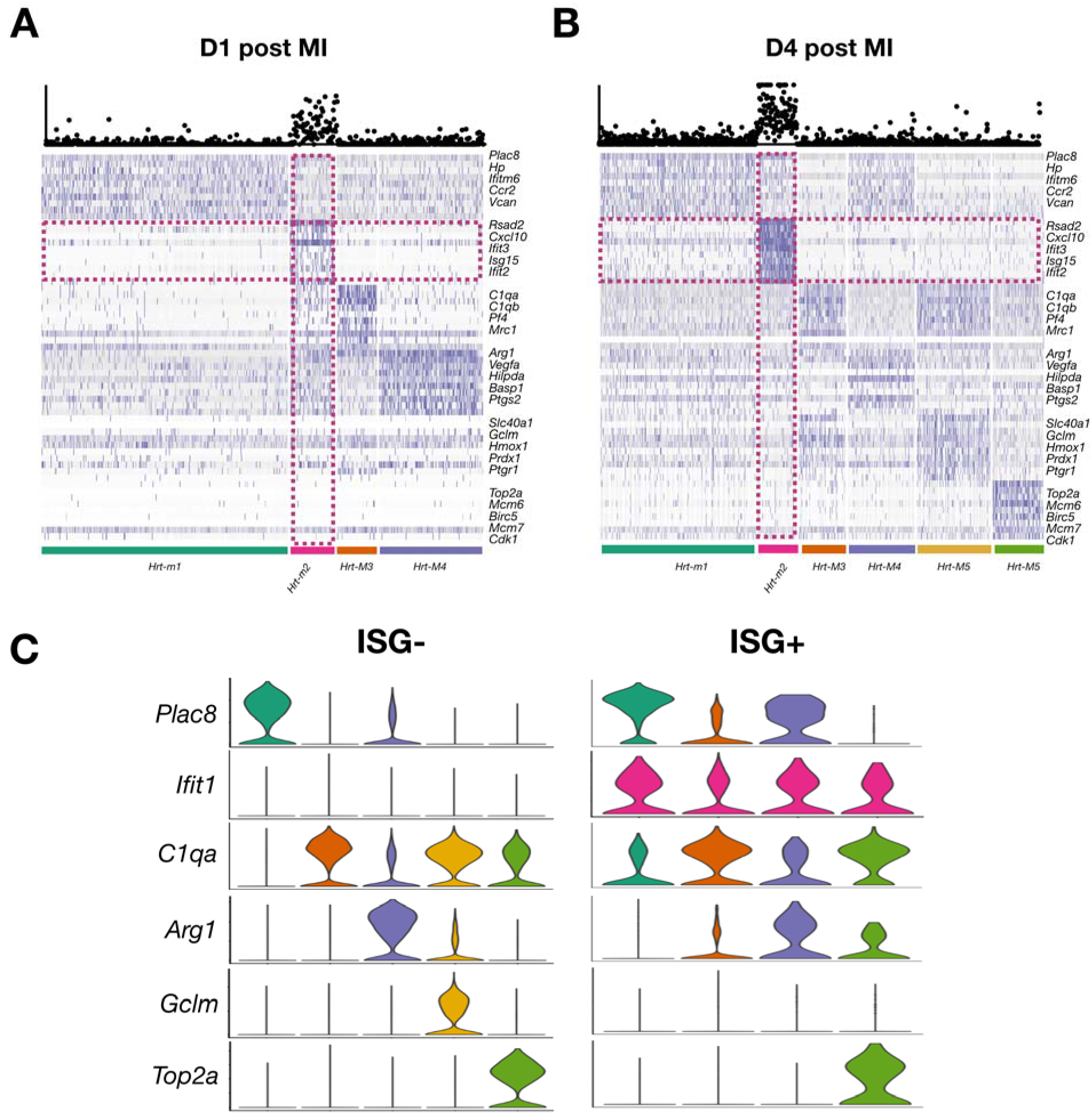
Single cell RNA seq. of WT cardiac macrophages on D1 and D4-post MI. (**A,B**) Representative heatmaps of cardiac macrophages from (**A**) D1 and (**B**) D4 after MI with ISG scores overlaid at the top. (**C**) Violin plots of bioinformatically isolated and subclustered ISG+ and ISG- macrophages. ISG+ cells can be subclustered into all ISG- subsets with the exception of the *Gclm* HI subset (gold).

**Figure S7.**
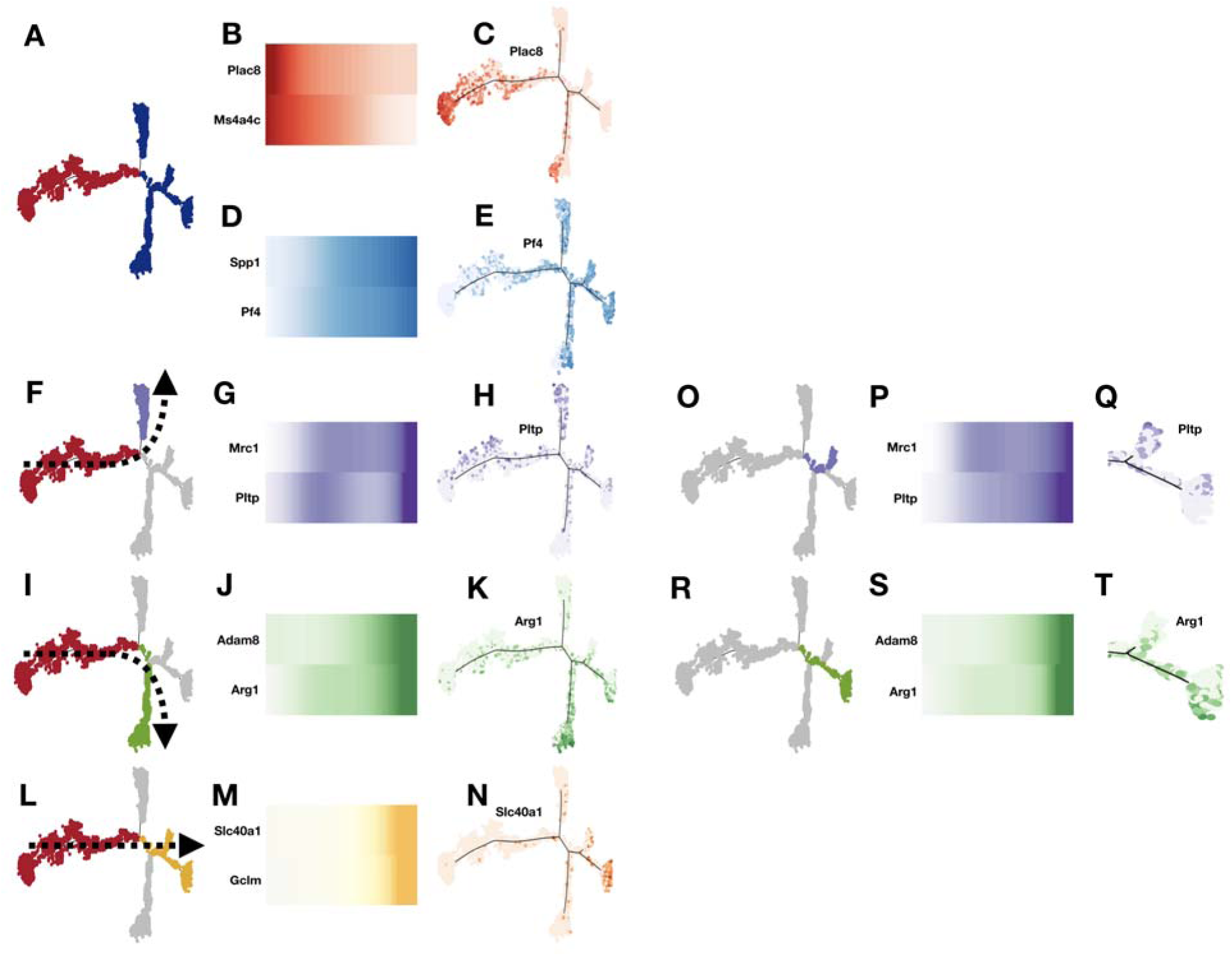
Pseudotime trajectory analysis of WT mice on day 4 after MI. **a)** Trajectory plot color coded by monocytes (red) and macrophages (blue). b) Heatmap of monocyte marker genes (*Plac8, Ms4a4c*) vs pseudotime. c) Trajectory plot highlighting monocyte marker gene, *Plac8*. d) Heatmap of monocyte marker genes (*Spp1, Pf4*) vs pseudotime. e) Trajectory plot highlighting macrophage marker gene, *Pf4*. f) Trajectory of monocyte to M1 macrophage differentiation. g) Heatmap of M1 macrophage marker genes (*Mrc1, Pltp*) vs pseudotime. h) Trajectory plot highlighting M1 macrophage marker gene, *Pltp*. i) Trajectory of monocyte to M2 macrophage differentiation. j) Heatmap of M2 macrophage marker genes (*Adam8, Arg1*) vs pseudotime. k) Trajectory plot highlighting M2 macrophage marker gene, *Arg1*. l) Trajectory of monocyte to Nrf2-dependent differentiation. m) Heatmap of Nrf2-macrophage marker genes (*Slc40a1, Gclm*) vs pseudotime. n) Trajectory plot highlighting Nrf2-macrophage marker gene, *Slc40a1*. o) Trajectory plot of M1-Nrf2-dependent differentiation. m) Heatmap of M1-Nrf2 marker genes (*Mrc1, Pltp*) vs pseudotime. n) Trajectory plot highlighting M1-Nrf2 marker gene, *Pltp*. r) Trajectory plot of M2-Nrf2-dependent differentiation. s) Heatmap of M2-Nrf2 marker genes (*Adam8, Arg1*) vs pseudotime. t) Trajectory plot highlighting M2-Nrf2 marker gene, *Arg1*.

**Figure S8.**
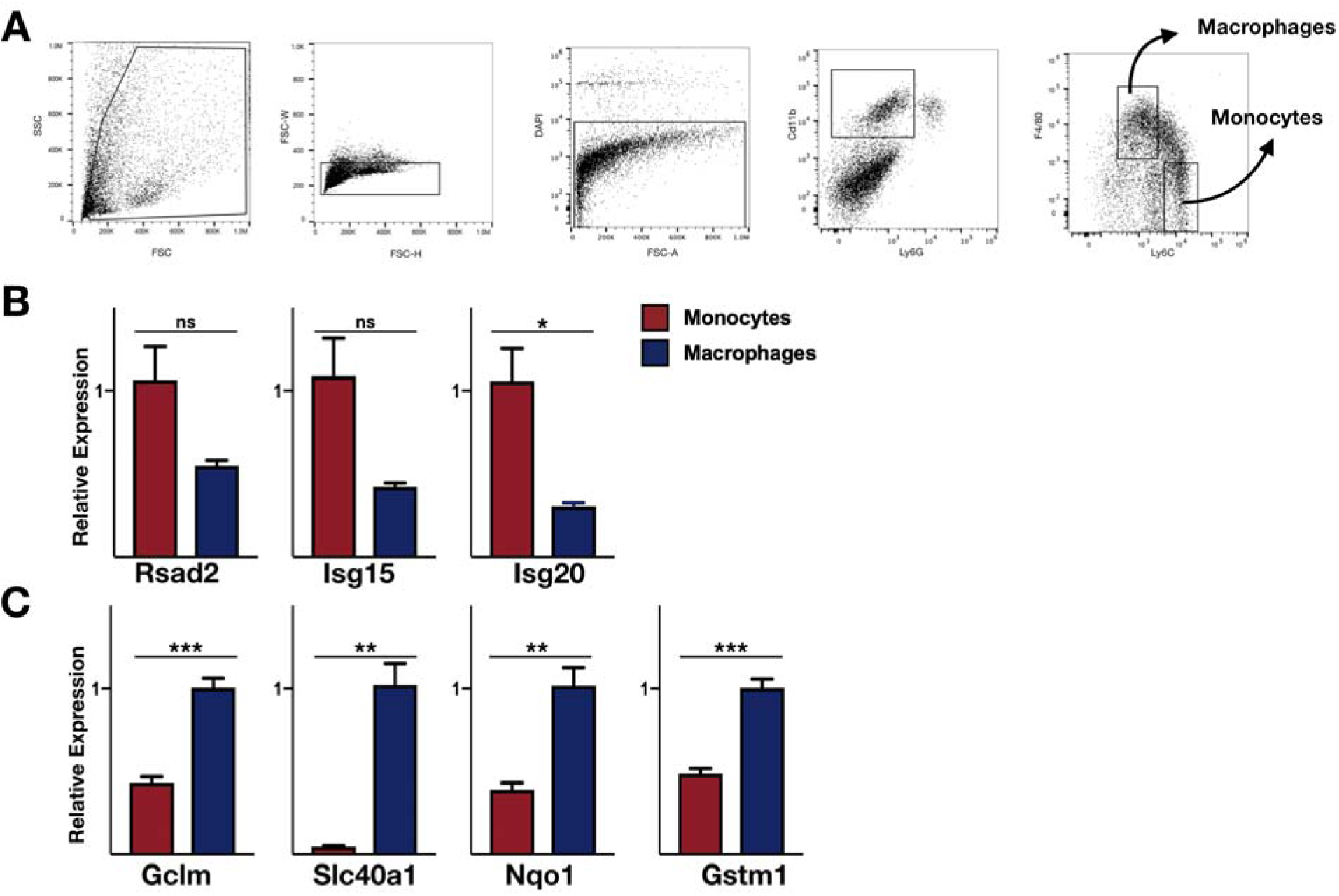
Quantitative PCR of sorted monocytes and macrophages from WT mice on day 4 after MI. a) Gating strategy for FACS sorting of infarct monocytes and macrophages. b) qPCR of ISGs (*Rsad2, Isg15, Isg20*) in sorted infarct monocytes and macrophages. c) qPCR of NSGs (*Gclm, Slc40a1, Nqo1, Gstm1*) in sorted infarct monocytes and macrophages.

**Figure S9.**
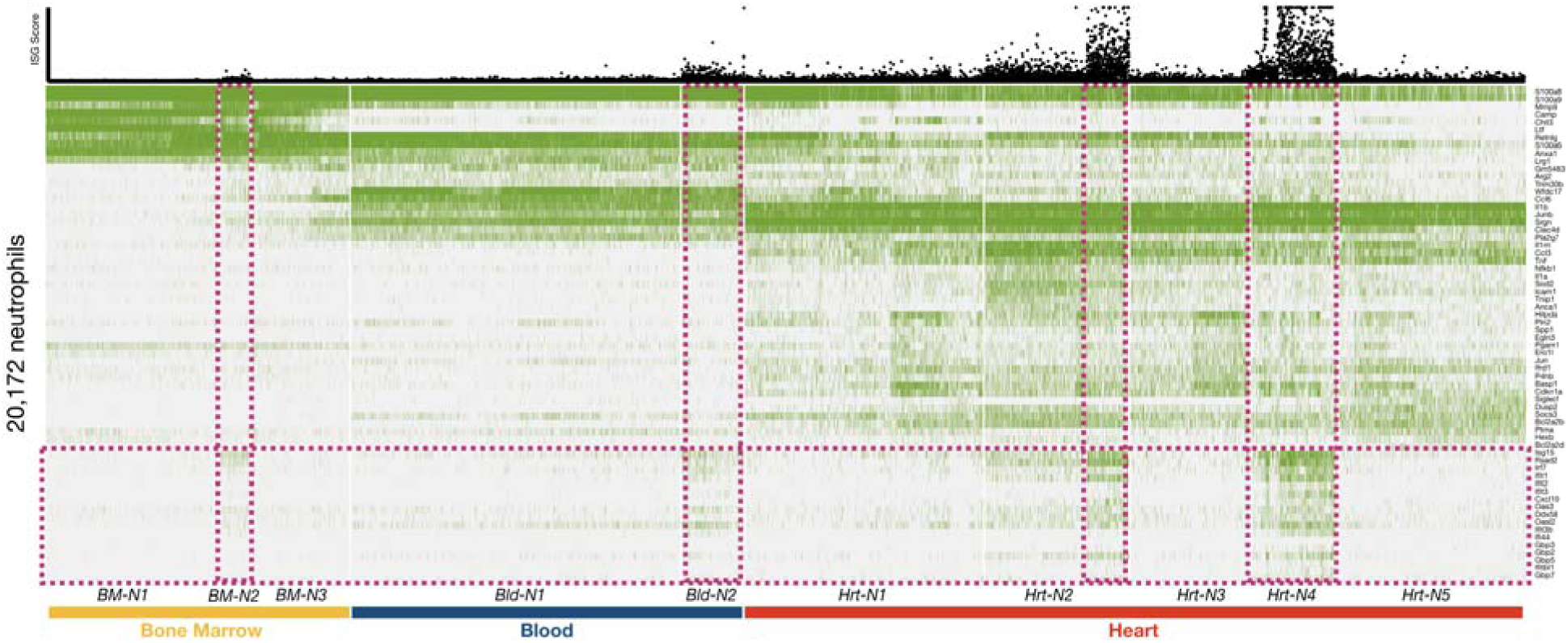
Interferon stimulation gradually increases from bone marrow through blood and to heart in neutrophils after MI. Composite heatmap of neutrophils (n = 20,172 transcriptomes) isolated on D1-D3 post-MI in mice from bone marrow, blood and heart. Maturation marker genes shown at top followed by blood markers, intracardiac differentiation markers, and ISGs at bottom, highlighted in pink.

**Figure S10.**
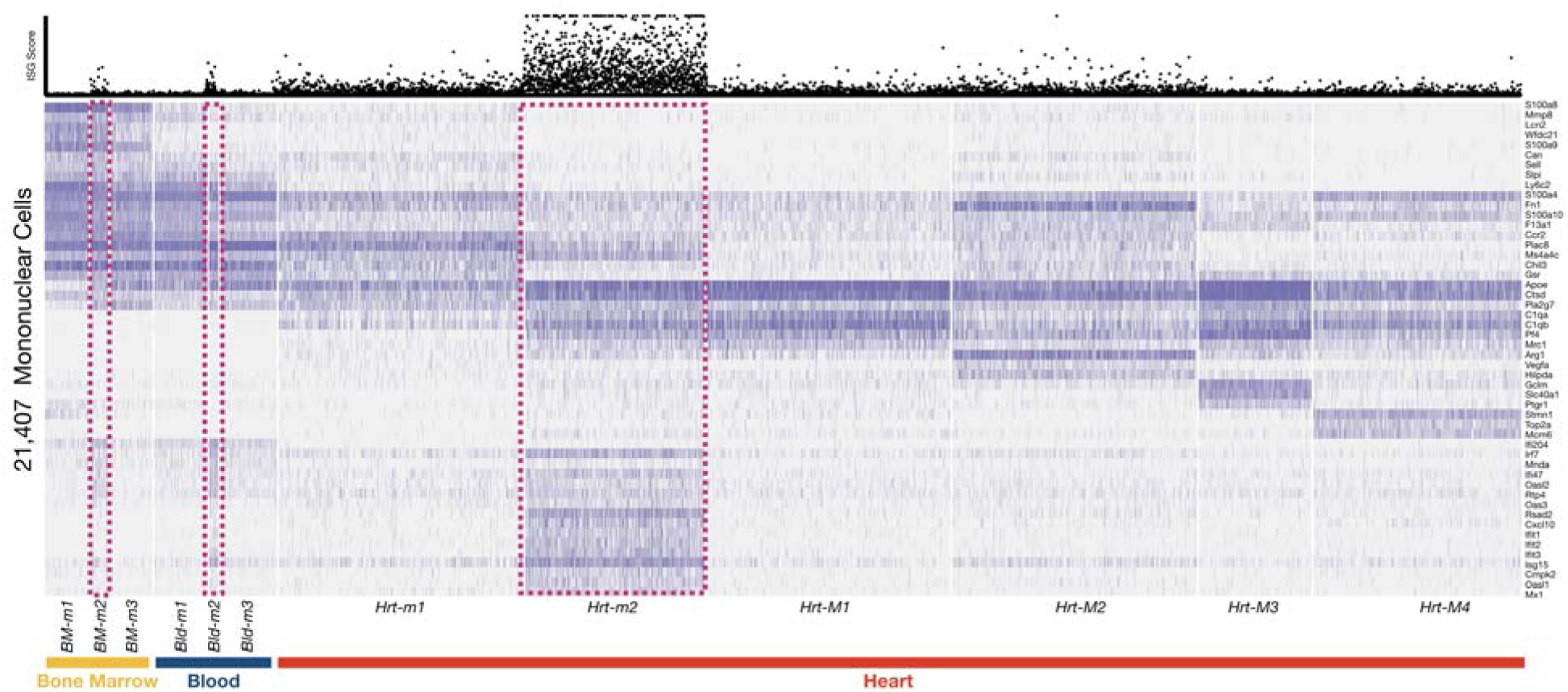
Interferon stimulation gradually increases from bone marrow through blood and to heart in monocytes after MI. Composite heatmap of monocytes (n = 21,407 transcriptomes) isolated on D1-D3 post-MI in mice from bone marrow, blood and heart. Maturation marker genes shown at top followed by blood markers, intracardiac differentiation markers, and ISGs at bottom, highlighted in pink.

## Materials and Methods

### Animals

Adult C57BL/6J (*WT,* stock 000664), Nrf2-deficient mice (stock 017009), Irf1-deficient mice C57BL/6NJ-Acod1<em1(IMPC)J>/J (stock 029340) mice were purchased from the Jackson Laboratory (stock 026554) or obtained from Fitzgerald lab after derivation from cryopreserved embryos obtained from the European Conditional Mouse Mutagenesis Program (EUCOMM)). IRF3^-/-^ mice^40^ were a generous gift from Tadatsugu Taniguchi and provided by Michael Diamond. IFNAR knockout mice (*IFNAR^-/-^*) were originally from J. Sprent and were backcrossed for 12 generations at the University of Massachusetts Medical School and provided by Kate Fitzgerald. Genotyping was performed in-house using methods recommended by Jackson Laboratory, or by Transnetyx. All experiments were performed with 10 to 14-week-old animals and were carried out using age and gender matched groups without randomization. All mice were maintained in a pathogen-free environment of the UC San Diego or Massachusetts General Hospital animal facilities, and all animal experiments were approved by the Subcommittee on Animal Research Care at UC San Diego or Massachusetts General Hospital.

### Myocardial Infarction and IFNAR Ab Treatment

Mice were intubated and ventilated with 2% isoflurane. After exposing the heart via thoracotomy at the fourth left intercostal space, the left coronary artery was permanently ligated with an 8-0 nylon monofilament suture. The thorax was closed with a 5-0 suture. Mice were treated with buprenorphine for analgesia on the day of surgery and twice daily thereafter for 72 hours. For cardioprotection experiments, mice were treated with two intraperitoneal doses of 500 μg of MAR1-5A3 IFNAR neutralizing antibody (BioXCell) at 8-12 hours and at 48 hours after permanent coronary ligation. Surgeries were performed in a blinded fashion unless genotype was obviated by unavoidable ascertainment of features such as coat color.

### Tissue Processing

Bone marrow cells were collected by flushing femurs with ice-cold PBC. The resulting solution was filtered through a 40 µm nylon mesh and treated with red blood cell (RBC) lysis (BioLegend). Blood was collected by cardiac puncture. The cellular fraction was collected into EDTA-containg tubse (Sgima), and erythrocytes were eliminated using RBC lysis. Heart tissue was collected by incising the right atrium and perfusing 10 mL of ice-cold saline into the left ventricular apex. Heart tissue was then removed and used for one of several assays detailed below. To obtain single cell suspensions for surface immunostaining, flow cytometric analysis, FACS sorting, or single cell RNA-Seq, hearts were enzymatically digested for 1 hour under continuous agitation in 450 U/ml collagenase I, 125 U/ml collagenase XI, 60 U/ml DNase I, and 60 U/ml hyaluronidase (Sigma) for 1 hour at 37°C, and filtered through a 40 µm nylon mesh in FACS buffer (DPBS with 2.5% bovine serum albumin) for enumeration by flow cytometry. For single cell RNA-Seq, the enzymatic digestion was limited to 45 minutes.

### Flow cytometry and Cell sorting

Isolated cells from enzymatically digested hearts were stained at 4°C in FACS buffer with DAPI to exclude dead cells, Ter119 (BioLegend, clone TER-119) to remove unlysed red blood cells, and mouse hematopoietic lineage markers directed against B220 (BioLegend, clone RA3-6B2), CD49b (BioLegend, clone DX5), CD90.2 (BioLegend, clone 53-2.1), NK1.1 (BioLegend, clone PK136). Secondary staining of leukocyte subsets was performed using CD45.2 (BioLegend, clone 104), CD11b (BioLegend, clone M1/70), Ly6G (BioLegend, clone 1A8), F4/80 (Biolegend, clone BM8) and/or Ly6C (BioLegend, clone HK1.4 or BD Bioscience, clone AL-21). Neutrophils were identified as (DAPI/B220/CD49b/CD90.2/NK1.1/Ter119)^low^ (CD45.2/CD11b/Ly6G)^high^. Monocytes were identified as (DAPI/B220/CD49b/CD90.2/Ly6G/NK1.1/Ter119)^low^ (CD45.2/CD11b)^high^ and sub-classified as pro-inflammatory monocytes F4/80^low^Ly6C^high^ or macrophages F4/80^high^Ly6C^low/int^. Flow cytometry was performed on an LSRII (BD Biosciences), SONY SH800, or SONY MA900, and analyzed with FlowJo software (Tree Star). Single cells from specific populations were collected for subsequent bimolecular analysis by FACS-sorting using a FACSAria II cell sorter (BD Biosystems), SONY SH800, or SONY MA900.

### Quantitative real-time PCR (qPCR)

Total RNA was extracted from FACS sorted cells or cultured cells using the RNeasy Micro kit (Qiagen) according to the manufacturer’s protocol. First-strand cDNA was synthesized using the High-Capacity RNA-to-cDNA kit (Applied Biosystems) according to the manufacturer’s instructions. TaqMan gene expression assays (Applied Biosystems) were used to quantify target genes (*Gapdh*: Mm99999915_g1, *Rsad2*: Mm00491265_m1, *Isg15*: Mm01705338_s1, *Isg20*: Mm00469585_m1, *Gclm*: Mm00514996_m1, *Gstm1*: Mm00833915_g1, *Slc40a1*: Mm01254822_m1, *Nqo1*: Mm01253561_m1). Relative changes were normalized to *Gapdh* mRNA using the 2^-ΔΔ^ method.

### Single Cell RNA-Seq

Single cell RNA-Seq was performed by microfluidic droplet-based encapsulation, barcoding, and library preparation, (inDrop and 10X genomics, see table 1 for sample level detail) as previously described (Engblom et al.). Paired end sequencing was performed on an Illumina Hiseq 2500 and Hiseq 4000 instrument. Low level analysis, including demultiplexing, mapping to a reference transcriptome (Ensembl Release 85 - GRCm38.p5), and eliminating redundant UMIs, was performed according to custom inDrops software (URL: https://github.com/indrops/indrops) [accessed April, 2017] or with CellRanger pipeline for 10X samples.

### Single-cell RNA-seq Data Quality Control, Normalization and Integration

To account for variations in sequencing depth, total transcript count for each cell was scaled to 10,000 molecules, and raw counts for each gene were normalized to the total transcript count associated with that cell and then natural log transformed. Cells with between 200 and 2,500 uniquely expressed genes and < 5% mitochondrial counts were retained for further analysis. Highly variable genes across individual datasets were identified with the *FindVariableFeatures* method from the Seurat R package (version 3.0) by selecting 2,000 genes with the highest feature variance after variance-stabilizing transformation.

Integration of multiple single-cell RNA-seq datasets was subsequently performed in Seurat to enable harmonized clustering and downstream comparative analyses across conditions(Butler et al., 2018; Qiu et al., 2017b; Stuart et al., 2019). Anchoring cell pairs between datasets were identified by Canonical Correlation Analysis (CCA) and the mutual nearest neighbors (MNN) method using the Seurat *FindIntegrationAnchors* function.

### Dimensional Reduction, Unsupervised Clustering, Sub-clustering

After scaling and centering expression values for each variable gene, linear dimensionality reduction was performed on integrated data using principal component analysis (PCA). Clustering was performed using the shared nearest neighbor (SNN) clustering algorithm with the Louvain method for modularity optimization, as implemented in the Seurat *FindNeighbors* and *FindClusters* functions. To visualize data in two-dimensional space, Uniform Manifold Approximation and Projection (UMAP) dimensional reduction was performed. Differentially expressed genes between clusters were determined using a Wilcoxon Rank Sum test.

As the highly-variable differentially expressed genes (DEGs) which drive unsupervised clustering are primarily composed of cell-type specific marker genes, we utilized a sub-clustering technique to examine cell-state heterogeneity within cell types in an unbiased manner. This involved serially subsetting particular cell-type clusters, identifying a new set of DEGs within the subset, and re-clustering the subset based on the newly determined DEGs.

### Pseudotime

Monocle (v3) package in R was utilized to order cells in pseudotime based on transitional expression patterns within the DEGs (Qiu et al., 2017a; Qiu et al., 2017b; Trapnell et al., 2014). Pseudotime trajectories were used to visualize the relationship between cell populations and pseudotime heatmaps were used to determine the dynamics of gene expression patterns in pseudotime. Only genes expressed in greater than 5% of cells were considered for dimensional reduction and differential analysis. For dimensional reduction, the number of dimensions was serially adjusted until branching local minimums were identified. Branching expression analysis modeling was implemented to determine branch dependent gene expression.

### ISG and NSG scores

ISG scores were measured as the sum of the raw reads for ten of the top ISGS: *Rsad2*, *Ifit2*, *Ifit3*, *Cmpk2*, *Cxcl10*, *Irf7*, *Isg15*, *Oasl1*, *Mx1*, and *Usp18*. NSG scores were measured as the sum of the raw reads for ten of the top NSGs: *Gclm*, *Slc40a1*, *Gstm1*, *Rnf128*, *Ptgr1*, *Treml4*, *Igf1*, *Cat*, *Ednrb*, and *Cd36*. Both ISG and NSG scores were log-normalized to reads per cell. ISG scores were thresholded to facilitate statistical comparisons between steady state and injured samples.

### Bone Marrow Derived Macrophage Stimulation

For *in vitro* stimulation with cyclic-di-GMP (Invivogen) (10ug/mL of media) and 4-Octyl-Itaconate (Tocris Bioscience) (100uM) was added to each well of a 6-well plate. After 24 hours of stimulation, RNA was extracted and quantified by qPCR as above.

### Statistics

Statistical analysis was performed using GraphPad Prism software. All data are represented as mean values +/- standard error of mean (S.E.M.) unless indicated otherwise. A statistical method was not used to predetermine sample size. For comparisons of the qPCR data and ISG/NSG scores a 2-tailed t-test with Welch’s correction was to determine statistical significance. All analyses were unpaired. *P* values are indicated by *P* values less than 0.05 were considered significant and are indicated by asterisks as follows: *p<0.05, **p<0.01, ***p<0.001, ****p<0.0001.

